# E2-Regulated Transcriptome Complexity Revealed by Long-Read Direct RNA Sequencing: From Isoform Discovery to Truncated Proteins

**DOI:** 10.1101/2025.07.31.667732

**Authors:** Didem Naz Dioken, Ibrahim Ozgul, Irem Yilmazbilek, Elanur Almeric, Irem Cemile Eroglu, Mustafa Cicek, Utku Cem Yilmaz, Deniz Karagozoglu, Esra Cicek, Ezgi Karaca, Tolga Can, Ayse Elif Erson-Bensan

## Abstract

Estrogen receptor alpha (ERα)-positive (ER+) breast cancers are driven by 17β-estradiol (E2) binding to ERα, which transcriptionally regulates downstream target genes. Although microarrays and conventional RNA sequencing have identified E2 target genes, pre-designed probes and short read lengths are limited in accurately capturing complex transcript structures. Long-read RNA sequencing offers a solution by spanning entire transcripts, providing a more complete view of the transcriptome. Here, we employed nanopore long-read direct RNA sequencing (DRS) complemented with 3’-end sequencing, in vitro experiments, and deep learning-based protein modeling to explore the intricate landscape of E2-responsive transcriptome and protein level implications. Our analysis revealed a range of E2-responsive non-coding and coding isoforms, including intronically polyadenylated (IPA) mRNAs. One of these IPA isoforms was detected for TLE1 (Transducin-like enhancer protein 1), which positively assists ERα-chromatin interactions for a subset of E2 target genes. The IPA isoform produces a C-terminus truncated protein, lacking the WDR interaction domain, but retains dimerization/tetramerization capacity through its intact N-terminus Q-domain. Structural modeling and protein-based assays confirmed the truncated protein’s dimerization potential and nuclear localization. Functional assays showed that overexpression of truncated TLE1 reduces the E2-induced upregulation of *GREB1*, an E2-responsive gene, thereby disrupting transcriptional regulation. Importantly, a lower IPA isoform ratio is associated with worse survival in ER+ patients, highlighting clinical relevance. Our study uncovers new layers of complexity in the E2-regulated transcriptome, providing insights into truncated proteins. These findings contribute to a deeper understanding of gene regulation and may help the development of new therapeutic strategies.

## Introduction

17β-estradiol (E2), the most potent circulating estrogen, plays a central role in regulating transcriptional responses that orchestrate diverse cellular processes essential for cell proliferation, differentiation, and survival. E2 primarily exerts its effects by binding to estrogen receptors (ER), particularly ERα, which activates the transcription of a wide range of genes [1]. ERα binds to over 10,000 genomic sites, recruiting transcription factors and co-regulators responsible for post-translational modifications of histones. These interactions and modifications influence the activity of the RNA polymerase II (RNAPol-II) transcriptional machinery, leading to the dynamic regulation of the transcriptome of E2-responsive cells [2].

Over the past few decades, transcriptomic analyses—primarily through microarrays and RNA sequencing (RNA-seq)—have substantially expanded our understanding of E2-regulated gene expression [1], [3]. These findings were pivotal in understanding molecular mechanisms underlying ER+ breast cancers and therapy responses.

RNA-seq remains the gold-standard technique for exploring transcriptomes with remarkable depth and precision. However, conventional short-read RNA sequencing methods have limitations, particularly in accurately capturing complex transcript structures and RNA modifications. Emerging long-read direct RNA sequencing (DRS) technologies offer a groundbreaking alternative by preserving transcript integrity and enabling the direct detection of diverse RNA species. This approach provides a more accurate representation of transcript isoforms, including those low in abundance or resulting from alternative splicing. By capturing this complexity, DRS offers new insights into transcriptome architecture, enhancing our understanding of how cells regulate responses to diverse physiological conditions, from yeast meiosis to stem cell development and diseases [4]. Therefore, a comprehensive understanding of transcriptome complexity is crucial for elucidating how RNA regulates protein function in normal and pathological states.

In this study, we used Oxford Nanopore Technologies (ONT) DRS, 3′-sequencing, protein structure modeling, and experimental validation to map a more detailed architecture of the E2-regulated transcriptome and investigate protein-level implications. Our findings revealed a complex landscape of non-coding and coding RNA isoforms, including intronically polyadenylated (IPA) transcripts from E2-responsive long genes, including *GREB1* and *TLE1*. We present experimental and computational evidence demonstrating the protein-level consequences for the C-terminally truncated TLE1 isoform, focusing on the lack of critical protein interaction regions. Notably, overexpression of the truncated TLE1 protein resulted in a reduced E2-driven upregulation of *GREB1*, indicating a disruption in the transcriptional regulatory network. Clinically, we show that a lower isoform ratio for TLE1-IPA is associated with poorer survival outcomes in ER+ breast cancer patients.

By highlighting E2-regulated IPA isoforms, our study sheds light on previously underexplored aspects of the E2-responsive transcriptome, suggesting significant functional implications at the protein level that may impact both breast cancer progression and potentially endocrine therapy response.

## Methods

### Cell lines and treatments

MCF7 cells were cultured in a phenol red-free starvation medium containing 10% dextran-coated charcoal-stripped FBS, 1% P/S, and 1% L-glutamine for 72 hours. After hormone deprivation (HD) for 72 hours, cells were treated with 100 nM 17 β-Estradiol (E2, Sigma-Aldrich, E2257) or ethanol (vehicle control) for 45 minutes, 3 hours, and 12 hours. RNA isolation, quantification, and cDNA synthesis were conducted according to MIQE guidelines [5]. Total RNA was isolated using the High Pure RNA Isolation Kit (Roche Applied Science) and incubated with DNase I (Roche Applied Science). cDNAs were synthesized using the RevertAid First Strand cDNA Synthesis Kit (Thermo Fisher Scientific). Cells were treated with actinomycin D (Tocris Bioscience) (10 µg/mL) to determine mRNA half-lives [6].

### QuantSeq 3’-Sequencing (3’-seq)

QuantSeq 3’-mRNA-Seq kit for Ion Torrent was used (Lexogen, CAT012.24A). The quality of RNA samples (RIN> 7) and the library were assessed by Agilent RNA 6000 Nano kit. IonXpress System kit (ThermoFisher) on the Ion GeneStudio S5 TM (ThermoFisher) and single-end run mode were used. Trim Galore was used for adapter removal and trimming (Krueger F., 2021, GitHub repository). The raw reads were aligned to the reference human genome GRCh37/hg19 using the TMAP aligner’s Map1 function and the BWA short-read algorithm. The resulting aligned BAM files were sorted using SAMtools and converted to bedgraph format using the BEDtools genomecov [7]. Bedgraph files with read counts were processed by MACS2 using the narrow peak option to identify QuantSeq peak boundaries [8]. Significant peaks at each time point and in the control group were identified with an FDR cutoff of 0.01. Peak boundaries were used to compute normalized read counts as Transcripts Per Million (TPM). No-polyA signal (or variants) [9], [10] harboring transcript ends (−60, +10) were removed from further analysis. polyA signal search parameters of the PolyASite 2.0 tool were used [11].

### Direct RNA sequencing (DRS)

MinION Mk1C and direct RNA sequencing protocols were used as suggested by Oxford Nanopore Technologies (ONT) (Direct RNA Sequencing Kit [SQK-RNA002] and Flow Cell [FLO-MIN106D]). Raw nanopore reads were aligned to the human genome GRCh37/hg19 and corrected using known splice junctions to increase the confidence in the generated isoform set. FLAIR was used for transcript identification and quantification [12]. For differential expression analysis, the “Diff_iso” module was used (fc > 1.5; fc<0.66). Transcripts with a minimum of 10 TPM in at least one of the E2 treatment time points were selected for further analysis. The transcript end locations of DRS reads were mapped to GRCh37. A custom Python script was used to overlap 3’seq reads (≥3 TPM) to DRS reads.

### Gene Ontology Analysis

For non-coding RNA enrichment analysis, we used the RNAenrich tool, which leverages experimentally validated RNA-target interactions to determine enriched biological processes and pathways [13]. Gene set enrichment analysis (GSEA) was performed using GSEA software (version 4.2.1) and pre-ranked analysis [14]. Public gene sets from the Broad Institute were downloaded using gene symbols. The number of permutations was set to 1000 and weighted enrichment statistics were used. Gene set enrichment analysis for ES and IR genes was performed by g:Profiler (version *e111_eg58_p18_f463989d*, database updated on 25/01/2024).

### RT-qPCR

MIQE guidelines were followed for RT-qPCRs [5]. RNA isolation, cDNA synthesis and RT-qPCR were performed as described. Ssoadvanced Universal SYBR Green Supermix (BioRad, CA, USA, 1725271) and CFX Connect Real-Time PCR Detection System (BioRad, 1855201) were used. Fold changes were normalized against the *RPLP0* reference gene. For relative quantification, the reaction efficiency incorporated ^ΔΔ^Ct formula was used [15]. Rapid amplification of cDNA ends (RACE)-specific cDNA synthesis was performed as described [6].

### In silico expression analysis

Single-cell RNA sequencing data from breast cancer patients (GSE75688) [16] was analyzed to detect intronic transcripts. A custom script was developed to identify samples containing intronic transcripts from BAM files. These transcripts were subsequently validated and visualized using Integrative Genomics Viewer (IGV). For bulk RNA sequencing (RNA-Seq) data, we utilized the TCGA-TARGET-GTEx project to compare the expression levels of target genes in normal tissues and tumors. RNA sequencing data for the Genotype Tissue Expression project (GTEx; https://gtexportal.org/home/) and The Cancer Genome Atlas (TCGA; https://portal.gdc.cancer.gov) was accessed via the Xena Toil RNA-Seq Recompute Compendium. GSE32474 [17], GSE78167 [18], and GSE27473 [19] were used for expression analysis.

### Patient survival analysis

Expression data for ENST00000376463.2 (IPA) and ENST00000376499.7 (FL) were downloaded from the TCGA/TARGET/GTEx study using the UCSC Xena browser. Clinical data of luminal A patients were downloaded from cBioPortal. IPA/FL ratios were calculated. Patients with overall survival times over 5 years or where both survival time and event status were zero were excluded. Patients were then stratified into high- and low-ratio groups using the best cutoff value that maximized the difference in survival. Kaplan-Meier survival curves show overall survival between the groups, and log-rank tests were performed to assess statistical significance. Microarray data of the KM plotter was used. The ratio of 228284_at (IPA) to 203221_at (FL) probe set expression values were determined for each patient. Patients were stratified into high- and low-ratio groups based on the optimal cutoff value to separate survival outcomes. The follow-up threshold was adjusted to 5 years.

### Truncated protein expression

The Coding Potential Calculator (CPC2) [20] was used to evaluate the truncated proteins’ reading frame. 3XFlag-GREB1-IPA and 3XFlag-TLE1-IPA were cloned in pcDNA3.1 (-) and transfected as described [6].NE-PER Nuclear and Cytoplasmic Extraction Kit was used (Thermo Scientific, 78835). For western blotting, a Flag antibody (Sigma F1804) was used. TUBA1 (α-Tubulin) antibody (Cell Signaling 2144) and HDAC antibody (68 kDa, Sc-81598 Santa Cruz) were used to confirm cytoplasmic and nuclear lysates.

### Immunocytochemistry (ICC)

Anti-Flag (Sigma F1804) primary, secondary antibodies (Abcam mouse IgG Alexa Fluor 488) and propidium iodide (Sigma, 81845) were used for ICC. Stained slides were visualized with an ANDOR AMH200 DSD2 Spinning Disc microscope with 100X (METU Biological Sciences).

### Protein Structures and Modelling

Protein amino acid sequences were taken from the UniProt Knowledgebase [21]. The crystal structures were retrieved from the Protein Data Bank [22]. The deep learning-based protein modeling tool AlphaFold2 (AF2) was used through the ColabFold platform [23], [24] to model truncated protein structures and their interactions. AF2 was used with an enhanced sampling protocol [25] on a local machine (Izmir Biomedicine and Genome Center, Karaca Lab). Per-residue confidence scores (pLDDT) (0−100), the predicted aligned error (PAE) rate for each residue, and the predicted structure accuracy (predicted TM (pTM)) (0-1) and interface predicted TM (ipTM) scores are used in the assessment of the AF2 models. PyMOL was used to visualize modeled structures (The PyMOL Molecular Graphics System, Version 2.0, Schrödinger, LLC, 1 December 2022). IBS 2.0 was used to illustrate the protein domain organizations [26].

## Results

Taking advantage of ONT DRS technology, we aimed to identify and characterize isoform diversity of E2-induced transcriptome. We sequenced native polyA+ RNA collected from hormone-deprived and three different time points (45 minutes, 3, and 12 hours) of E2 (100 nM) treated MCF7 cells, a well-established model for E2 responsive ER+ breast cancers. DRS generated around one million reads, with a minimum of 92% pass reads and a mean per-base quality score (PHRED) of >9.9/sample. Sequenced read length means were approximately 1.1 kb, whereas the maximum read lengths ranged from 15 kb to 49 kb across different samples (Table S1 for QC reports). We used FLAIR (Full-Length Alternative Isoform analysis of RNA) to identify high-confidence transcript structures [12]. Among the differentially expressed transcripts (Figure S1), most of the sequenced transcripts (94%) belonged to protein-coding genes, and the remaining were lncRNAs (4%) and other (anti-sense, processed pseudogenes) non-coding RNAs (ncRNAs) (2%) (Figure 1a). As anticipated, the DRS-sequenced transcripts were enriched for E2-responsive genes (Table S2), one of which was TFF*1,* a known E2-responsive gene (Figure S2). There was an initial burst of upregulated and downregulated genes at 45 minutes, and by 12 hours, the number of upregulated genes decreased. At 12 hours, there was a higher number of downregulated genes, a pattern in agreement with previous reports (Figure 1b) [2], [27].

**Figure 1.**
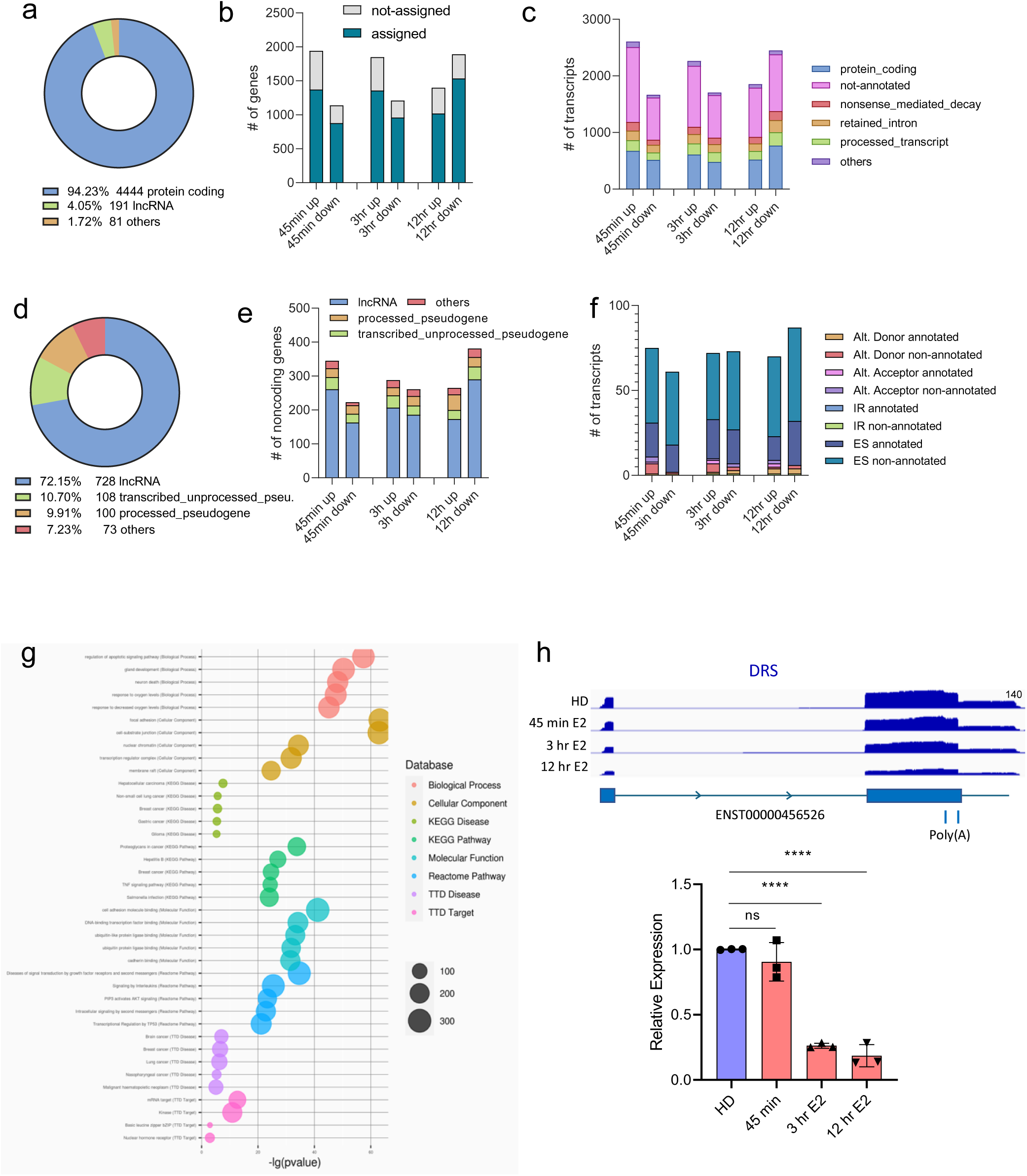
**a.** Differentially expressed gene categories (TPM>10 at any time point, fc>1.5, fc<0.6) compared with the hormone-deprived control, **b**. The number of upregulated (fc>1.5) and downregulated (fc < 0.6) genes at 45 min., 3 hrs., and 12 hrs. of E2 treatment. Blue bars represent genes assigned to ENSEMBL genes, while grey bars indicate reads that were not assigned a gene ID, **c.** Upregulated and downregulated transcripts at different time points. Transcripts are classified according to ENSEMBL Biotype, represented by distinct colors, **d**. Categories of differentially expressed non-coding transcripts, (TPM>2 at any time point), **e**. The number of upregulated and downregulated non-coding gene types, **f.** Types of splicing events for ncRNAs captured by FLAIR, **g.** RNAenrich analysis of differentially expressed ncRNA genes, Benjamini-Hochberg (FRD) adjustment, p<0.05), **h.** IGV visualization of DRS reads for lncGATA3-7 (top), RT-qPCR confirmation of lnc-GATA3-7 downregulation upon E2 stimulation compared with hormone-deprived (HD) MCF7 cells. Expression levels were normalized using *RPLP0* as the reference gene (n=3 independent E2 treatments, Statistical significance was evaluated by one-way ANOVA, with **** p < 0.0001, ns: not significant.).

Having successfully captured the expected gene expression profile following E2 stimulation, we next looked deeper into the transcriptome, leveraging the isoform discovery and characterization capabilities of DRS and the FLAIR algorithm [12]. Some of the sequenced E2-regulated transcripts were not assigned a gene code (ENSG) by FLAIR merely because there was no complete alignment with the canonical transcription start sites/ends. Hence, the “not-assigned/non-annotated” transcripts do not necessarily originate from novel genes (Figure 1b).

Likewise, when we investigated the ENSEMBL Biotype classification of E2-regulated transcripts (Figure 1c), the “non-annotated” category consisted of transcripts that were not identical to known isoforms; hence FLAIR did not assign an existing transcript code (ENST). The next group of E2-regulated transcripts belonged to the annotated protein-coding transcripts, processed transcripts (without an open reading frame), and nonsense-mediated decay transcripts with potentially in-frame termination codons 50 bp upstream or more of the final splice junction (Figure 1c).

We first looked into the non-coding component of the E2-regulated transcriptome to uncover new layers of transcriptome complexity.

### E2-regulated non-coding RNAs

The E2-regulated ncRNA transcripts consisted mostly of lncRNAs (n=728, 72%), transcribed unprocessed pseudogenes (n= 108, 11%), processed pseudogenes (n=100, 10%), and others (7%) (Figure 1d). Like the protein-coding genes, most ncRNAs were initially upregulated and the number of downregulated non-coding RNAs increased at 12 hours of E2 treatment (Figure 1e). The most common alternative processing for ncRNA transcripts was exon skipping (Figure 1f). This pattern agreed with recent reports on the widespread splice variants for lncRNAs [28].

These E2-regulated ncRNAs were significantly enriched for apoptotic signaling, gland development, and other cancer-relevant processes, determined by the RNAenrich tool, which utilizes experimentally validated RNA-target interactions (Figure 1g) [13].

At this point, we wanted to experimentally confirm one of the downregulated ncRNAs detected by DRS because lncRNA levels are generally comparably lower than mRNAs [29]. According to DRS, E2 treatment caused the downregulation of a spliced and 270 nucleotide long lncGATA3-7 (LNCipedia, ENST00000456526.1, ENSG00000223808.1). The significant downregulation of lncGATA3-7 upon E2 treatment was confirmed by RT-qPCR (Figure 1h). lncGATA3-7 was previously described as a luminal type lncRNA (LOL) [30], and its expression is low in non-ER+ breast cancer cells (Figure S3). Of note, lnc-GATA3-7 is uniquely expressed in only a few tissues, including the normal mammary tissue (Figure S3).

Exemplified by lncGATA3-7, the coordinated and dynamic expression of ncRNAs in response to E2 underscores their functionality during the E2 response. Deciphering the sequence isoforms of ncRNAs—whether generated by splicing or through alternative transcript start/stop sites—is crucial for understanding their functions, given their cell type-specific expression patterns and roles in various physiological processes [31].

We then looked into the diversity of transcript isoforms within protein-coding genes, beginning with the characterization of differential splicing patterns, followed by an analysis of mRNA 3’-ends.

### Spliced Isoforms

There was a dynamic upregulation of alternatively spliced annotated and non-annotated transcripts in 45 min., 3, or 12 hrs. E2 treated cells (Figure 2a). This landscape of E2-regulated transcripts was grouped into alternative donor (5’-ends of introns), alternative acceptor (3’-ends of introns), intron retention (IR), or exon skipping (ES) events. ES was the most common event for annotated and non-annotated isoforms (Figure 2a). IR was more common for the non-annotated isoforms, suggesting that DRS could identify uncharacterized isoforms (Figure S4). DRS was able to capture ES and IR transcripts for *CRB3* (Crumbs cell polarity complex component 3) and *HAGH* (Hydroxyacylglutathione hydrolase) (Figure 2b, 2c). CRB3 is an important regulator for breast cancer stemness, which is associated with tamoxifen resistance [32]. HAGH is involved in glutathione biosynthesis and detoxification of reactive and toxic 2-oxoaldehydes [33].

**Figure 2.**
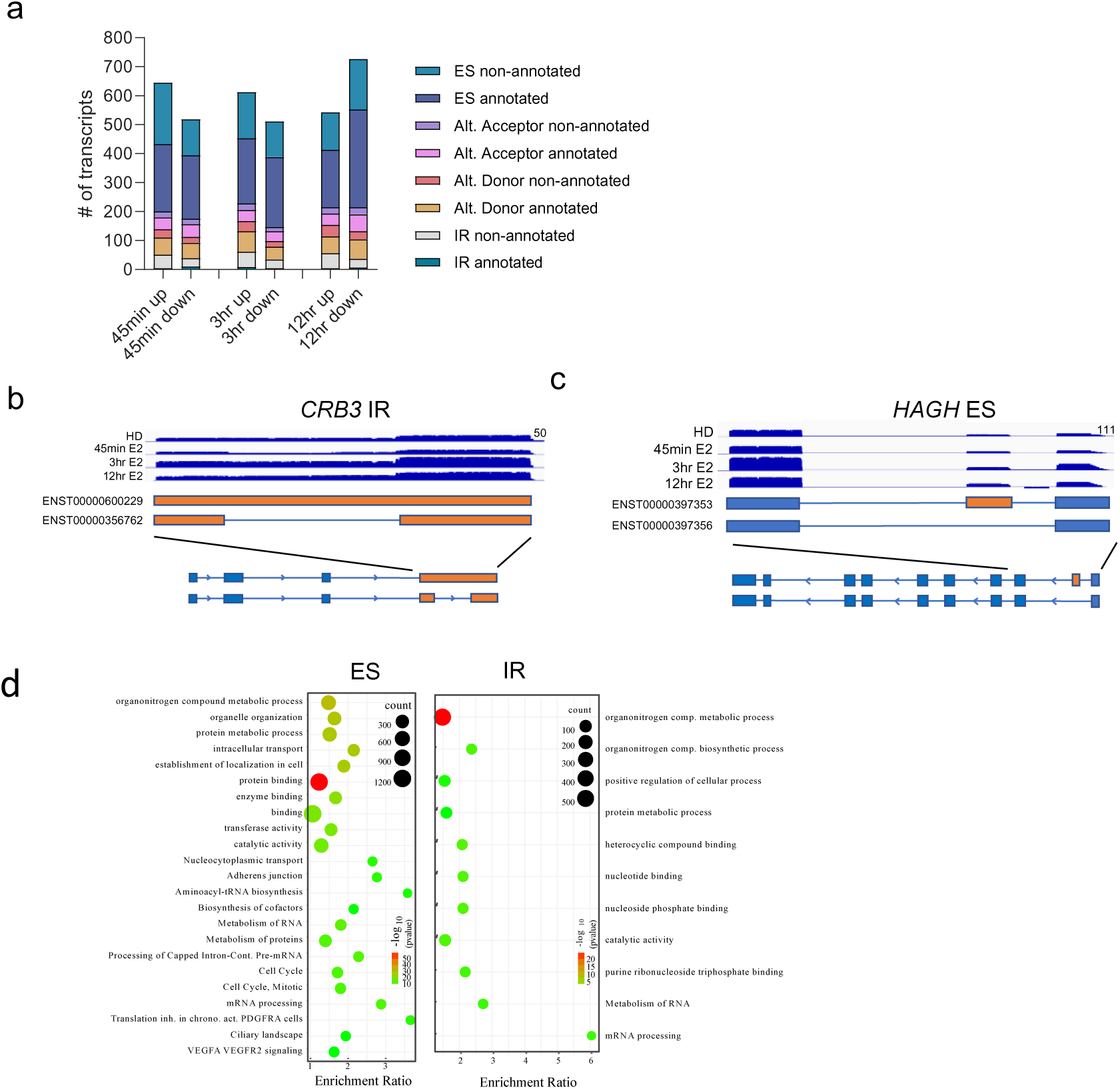
**a**. Number of alternative transcripts up or down-regulated upon E2 stimulation. Events are indicated in the key and as colored bars, **b**. IGV visualization of DRS reads capturing an isoform with intron retention (IR) for CRB3, **c.** IGV visualization of DRS reads capturing an isoform with exon skipping (ES) for HAGH, **d.** Gene ontology enrichment analysis for ES and IR groups. Enriched biological processes and molecular functions are shown as bubble plots. The x-axis represents the enrichment ratio, the y-axis lists the top enriched terms. The size of each bubble corresponds to the number of genes associated with the term, and the color represents the significance level (-log10 of the p-value).

Overall, the ES and IR cases were enriched for transcripts involved in the metabolic process (GO: BP), protein binding (GO: MF), and nucleotide binding (GO: MF) (Figure 2d). These cases may have various implications, depending on the inclusion or exclusion of alternate exon and intron sequences and the resulting alternative domain architectures.

### Transcript 3’-ends

To identify and confirm isoforms with different 3’-ends, we combined DRS and 3’-seq results. Both sequencing methods identified transcript 3’-ends mapping to the expected terminal exons, but they also detected unexpected reads mapping to intronic regions within the genome. To explore these intronic regions, we classified the differentially regulated (fc >1.5, TPM >10, or fc < 0.6, TPM> 10) transcripts based on their 3′-ends in the human genome. As a result, the transcript ends were organized into three categories: early intron (EI), mid-intron (MI), and terminal exon (TE).

The TE group includes all sequenced transcripts that terminate in the last exon, with various alternative splicing events (i.e., Alt 5’, Alt 3’, ES, and IR). 3054 transcripts that end in terminal exons were upregulated (TPM > 10 and Isoform Ratio > 10% at any time), including 1219 non-annotated transcripts. The 3’-seq had overlapping reads with 2075 DRS-captured transcript 3’- ends. For the EI and MI groups, DRS captured 413 intronically polyadenylated (IPA) and upregulated transcripts. Of these, 246 were not previously annotated. 3’Seq detected 188 of these transcript ends.

### EI and MI transcripts

A significant number of transcripts of the EI (n=110) and MI (n=240) groups were differentially expressed at 45 min. of E2 treatment compared to TE transcripts (n=2751) (Figure 3a). The upregulated EI transcripts at 45 min. of E2 treatment gradually decreased at later time points. Moreover, new EI and MI transcripts were upregulated at 3 and 12 hr. time points (Figure 3a). To begin understanding these IPA transcripts, we investigated gene and intron sizes. Both EI and MI transcripts originate from significantly longer genes than the TE genes (Figure 3b). The EI transcripts had the longest early intron (intron1 or intron 2) sizes. Both the EI and MI transcript 3’-ends (−60 to +10) had at least one of the common polyadenylation signals (AAUAAA) and/or its variants [9] (Figure 3c). We did not detect common poly(A) signals only in 10% of non-annotated EI transcripts, and in less than 5% of other transcripts. Overall, these suggested that most IPA transcripts originated from longer genes and longer early introns in the case of EI transcripts with conserved poly(A) signals.

**Figure 3.**
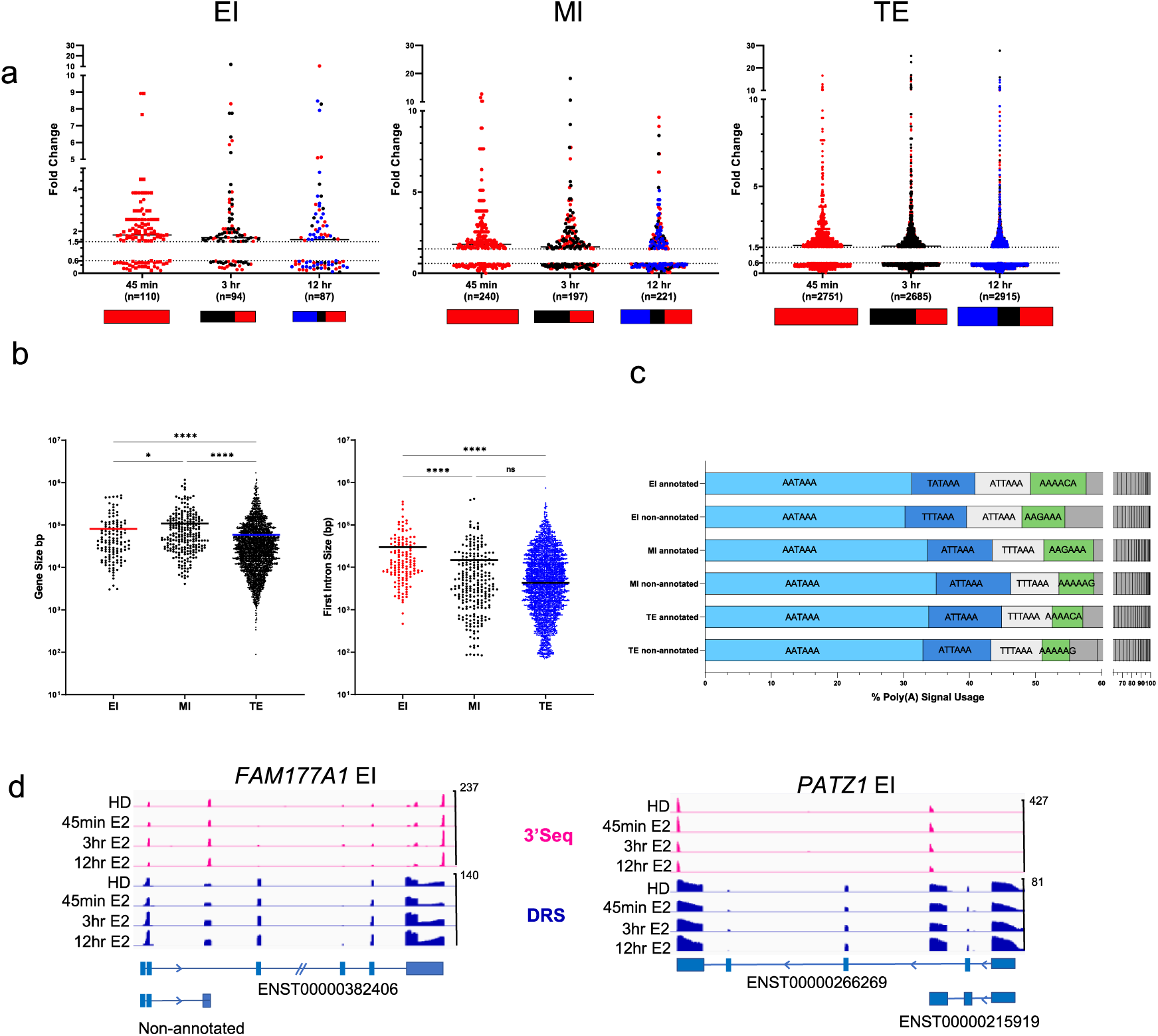
**a.** EI, MI, and TE group of upregulated (>1.5 fold) and downregulated (<0.6 fold) transcripts identified by DRS. Red, black, and blue dots/boxes indicate transcripts significantly altered at 45 min., 3, and 12 hrs., **b.** Gene size (left) and early intron (first or second introns) size (right) of upregulated EI, MI, and TE genes. Horizontal bars indicate the median values (Kruskal-Wallis, multiple pairwise comparisons, *p<0.05, ****p<0.0001, ns: not significant), **c.** One or more common polyadenylation signals detected for the annotated and non-annotated EI, MI, and TE genes, **d**. IGV display of DRS and 3’seq reads showing IPA and FL transcripts of *FAM177A1* and *PATZ1*.

For the EI group, IPA transcripts included both annotated and non-annotated transcripts that terminate within early introns, as exemplified by *FAM177A1* and *PATZ1*. The 3’-seq data also confirmed the presence of these intronically terminated transcript ends (Figure 3d).

In addition to these EI transcripts detected by DRS, we observed a distinct set of 3’-seq reads that were not captured by DRS. These early intronic 3’-seq peaks were initially eliminated from our analysis because they were near (−100 to +100) A-stretch sequences associated with SINE/Alu elements in early and long introns (Figure S5). We suspected these reads were due to internal oligo-dT priming from pre-mRNA templates. Still, we were able to confirm upregulated expression using random-hexamer cDNAs and RT-qPCR for two E2-responsive genes, *IGFBP4* and *TPD52L1* (Figure S6). Curiously, independent evidence from an Affymetrix probe set (242657_at) also shows widespread expression from the same intronic region of *IGFBP4* (Figure S7).

Interestingly, these 3’-seq reads from early intronic regions were sharply upregulated at 45 minutes, then gradually decreased, while their full-length transcripts showed a slower increase (Figure S8). Notably, at 45 minutes, these genes were only modestly upregulated (based on 3’UTR reads) compared with genes that only had terminal exon 3’seq reads (Figure S9). A similar expression pattern was observed in an independent RNA-seq dataset (GSE78167) of E2-treated MCF7 cells (Figure S9). Based on these results, we predict that these EI 3’-seq reads not captured by DRS may correspond to delayed processing of long pre-mRNAs. This delay could affect full-length transcript levels by hindering efficient intron removal—a reported cause of significant delays in gene expression [34]. Previous studies have also reported that internal priming during sequencing and cDNA synthesis can occur at intronic Alu elements. These observations are linked to delayed splicing and the presence of pre-mRNAs in total RNA isolates [35]. Our observations align with these reports, also supporting the view that delayed processing of long pre-mRNAs may contribute to the fine-tuning of gene expression. Further work will be interesting to fully understand the regulatory consequences of these delayed processing events and their broader implications for transcriptome dynamics.

Next, within the IPA isoforms, we focused on the MI group as they are likely to generate stable mRNAs translated into C-terminus truncated proteins. We selected MI group candidates that are involved in the E2 response. *GREB1* (Growth Regulating Estrogen Receptor Binding 1) is an E2-responsive gene [36] and *TLE1* (TLE family member 1) mediates estrogen receptor binding and transcriptional activity in breast cancer cells [37]. DRS and 3’seq revealed intronically terminated minor isoforms for both *GREB1* and *TLE1* (Figure 4a). DRS and 3’-seq showed the upregulation of an IPA minor isoform for *GREB1,* terminating within the seventh intron. The *TLE1* IPA minor isoform terminated within the sixth intron.

**Figure 4.**
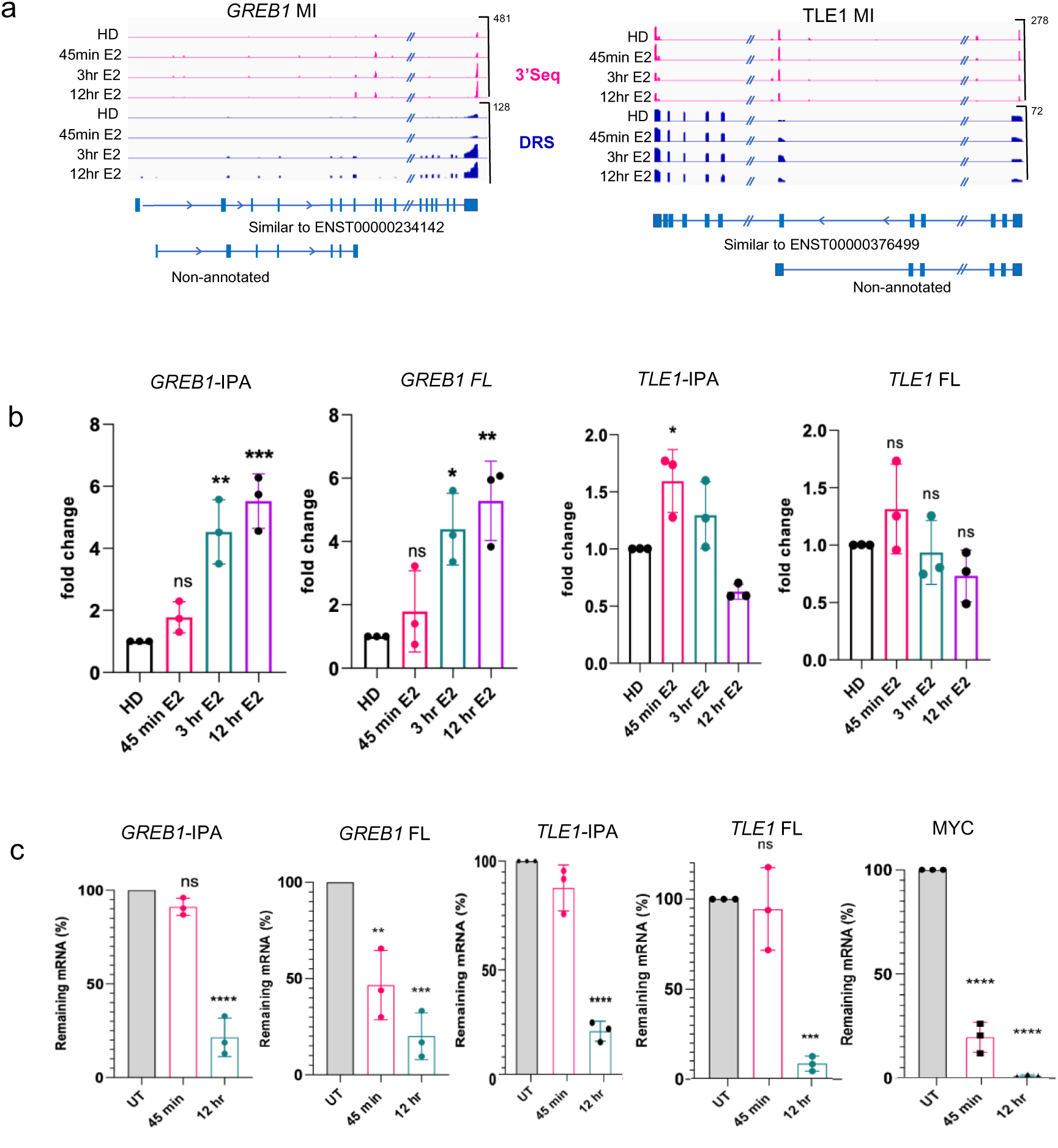
**a.** IGV display of 3’seq and DRS reads and transcripts defined by FLAIR. In addition to the full-length mRNAs, intronically terminated isoforms of GREB1 and TLE1 were detected, **b.** RT-qPCR for the expression of IPA and FL isoforms of *GREB1* and TLE1 upon E2 stimulation relative to hormone-deprived (HD) MCF7 cells. The data represent 3 independent E2 treatments. One-way ANOVA with Tukey’s multiple comparison test was used, ns: not significant, \**p* < 0.05, \*\**p* < 0.01, ***p<0.005, **c.** Stabilities of IPA and FL isoforms detected by actinomycin D (10 µg/ml) treatment. RT-qPCR results show remaining mRNA levels at 45 min and 12 hr. of actinomycin treatment. *MYC* is a short-lived mRNA and was used as a positive control for the treatment. The data represent the mean (SD) of 3 independent treatments. One-way ANOVA with Tukey’s multiple comparison test was used \*\*\**p* < 0.001, \*\*\*\**p* < 0.0001.

*GREB1* IPA isoform was gradually upregulated throughout the E2 treatments. The *TLE1* IPA isoform was initially upregulated but later downregulated upon E2 stimulation. The 3’-ends of both IPA isoforms were confirmed by 3’seq (Figure 4a). These IPA isoforms were not assigned ENST codes by FLAIR.

To begin characterizing these IPA isoforms, we first experimentally confirmed their expression using an independent set of E2-treated cells. Hormone-deprived (HD) MCF7 cells were treated with E2, and total RNA was isolated. RT-qPCR results showed a similar pattern of E2-regulated expression (Figure 4b). The 3’-end of the IPA isoforms was further confirmed by 3’RACE (Rapid amplification of cDNA ends) and sequencing of the transcript 3’-ends (Figure S10, S11). Both IPA isoforms had comparable mRNA half-lives with the full-length isoforms detected by blocking transcription and quantifying remaining RNA levels by RT-qPCR (Figure 4c). *MYC* mRNA has a significantly short half-life [38] and was used as a positive control in this experiment.

We sought additional evidence to confirm the presence of these intronically terminated isoforms for *GREB1* and *TLE1*. We looked into independent datasets from various sequencing platforms to examine the expression of IPA isoforms. Notably, both IPA isoforms were detected in single-cell RNA seq data of breast cancer cells (GSE75688) (Figure 5a). In addition, we found a microarray probe set to specifically recognize the *TLE1*-IPA isoform (228284_at), showing widespread expression in different cells (Figure 5b). Utilizing GEO2R, we detected decreased expression of the *TLE1*-IPA isoform (228284_at) and the full-length TLE1 (203221_at) in ERα knock-down (KD) MCF7 cells (Figure 5c). These results showed independent evidence of the intronically terminated isoform and its ERα dependency. However, there was no pre-designed microarray probe set to recognize the *GREB1*-IPA.

**Figure 5.**
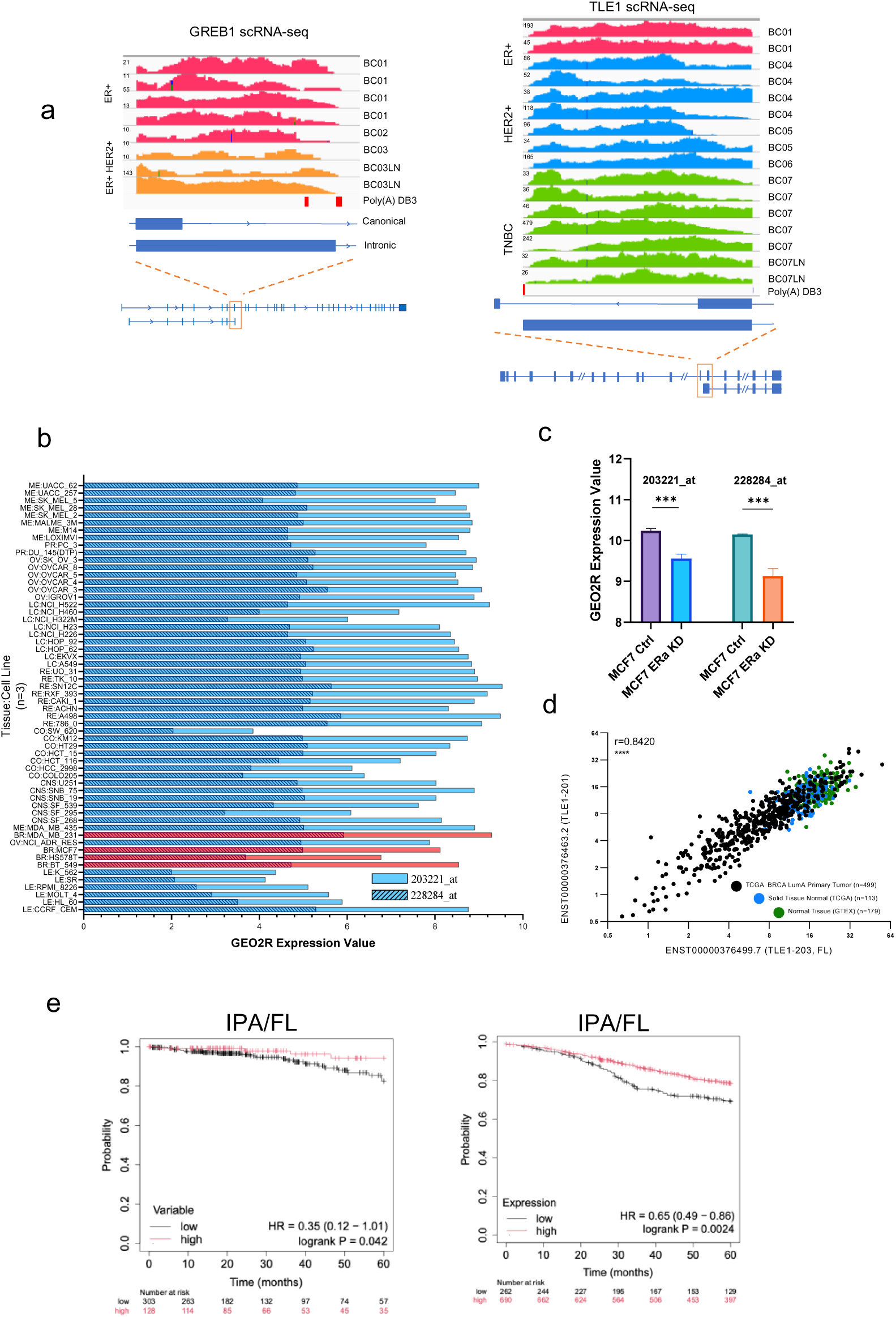
**a.** Analysis of single-cell RNA sequencing data of GSE75688 verified the expression of IPA isoforms of *TLE1* and *GREB1* in breast cancer cells representing subtypes, **b.** GEO2R expression data for *TLE1*-IPA (203221_at probe set) and FL isoforms (228284_at probe set) in NCI-60 Cancer Cell Line data (GSE32474, GPL570). Red bars indicate breast cancer cell lines, **c.** Probe sets specific to IPA (203221_at) and FL (228284_at) in control cells compared with ER knockdown (KD) MCF7 cells (GSE27473) (***p<0.001, students t-test), **d.** TCGA breast cancer dataset showing a positive correlation between the levels of the FL and IPA isoforms of *TLE1*, **e.** Kaplan-Meier survival curves comparing high- and low-ratio of TLE1 ENST00000376463.2/ ENST00000376499.7 (IPA/FL) TLE1 in TCGA luminal A breast cancers. High-ratio patients (red) showed better progression-free survival compared to low-ratio patients (black) (HR = 0.47, log-rank p = 0.033) (Left panel). Kaplan-Meier survival analysis of breast cancer microarray data from KM plotter, stratifying patients by the ratio of 228284_at (IPA) to 203221_at (FL) probe set expression. High-ratio group (red) exhibited improved survival compared to the low-ratio group (black) (HR = 0.65, log-rank p = 0.0024) (Right panel). Patient numbers and at-risk counts are displayed on the plot.

Next, we investigated GTEx (The Genotype-Tissue Expression) for normal breast and TCGA (The Cancer Genome Atlas) RNA-seq datasets for breast cancer patient samples. For *TLE1*, GTEx lists ENST00000376499.7 (TLE1-203, 4135 bp, 770 aa) as the major and full-length transcript in normal breast tissue. Of note, there is a second *TLE1* transcript (ENST00000376463.2), and it has an intronically terminated 3’-end, identical to the DRS-captured IPA isoform. This isoform is indicated in ENSEMBL as TLE1-201 (1115 bp) but is marked to lack EST evidence at the 5’end, and hence, the start of the coding sequence is not annotated (Figure S12). In contrast, the DRS-captured IPA isoform delineates the 5′ and 3′ ends of the transcript. Consequently, we regarded the DRS-captured IPA isoform as the complete version of TLE1-201 and investigated its expression pattern as an indicator of the IPA isoform’s presence in breast tumors. Analysis of the TCGA dataset revealed a positive correlation between the levels of the full-length *TLE1* transcript and the TLE1-201 isoform in breast cancers (Figure 5d). These results suggested that the intronically terminated isoform is not rare and co-exists with the full-length isoform. Moreover, a lower IPA/FL isoform ratio is associated with worse survival in ER+ breast cancer patients (luminal A) (Figure 5e), suggesting functional implications for the *TLE1* IPA isoform.

On the other hand, *GREB1* is not abundantly expressed in normal breast tissue (Figure S13) but is known to be upregulated by E2 [36].

### Structural Modeling

These findings prompted us to investigate the protein produced from the IPA isoforms. The DRS-captured *TLE1* IPA isoform exhibits a similar coding potential to the full-length isoform (Figure 6a). This isoform codes for 200 amino acids, in contrast to the 770 amino acids translated by the full-length TLE1 (Figure 6b). The full-length TLE1 consists of an N-terminal glutamine-enriched (Q) domain, a Gly-Pro-enriched (GP) domain, a CcN cysteine-rich domain, a Ser-Pro-enriched (SP) and tryptophan-aspartic acid repeat region (WDR) [39]. In the TLE-IPA isoform, eight new amino acids are introduced after the 192^nd^ amino acid before a STOP codon is encountered. The isoform loses all full-length domains except for the Q-domain, linked to the dimerization and tetramerization of TLE1 [39] and the GP domain (131-192 aa) (Figure 6b).

**Figure 6.**
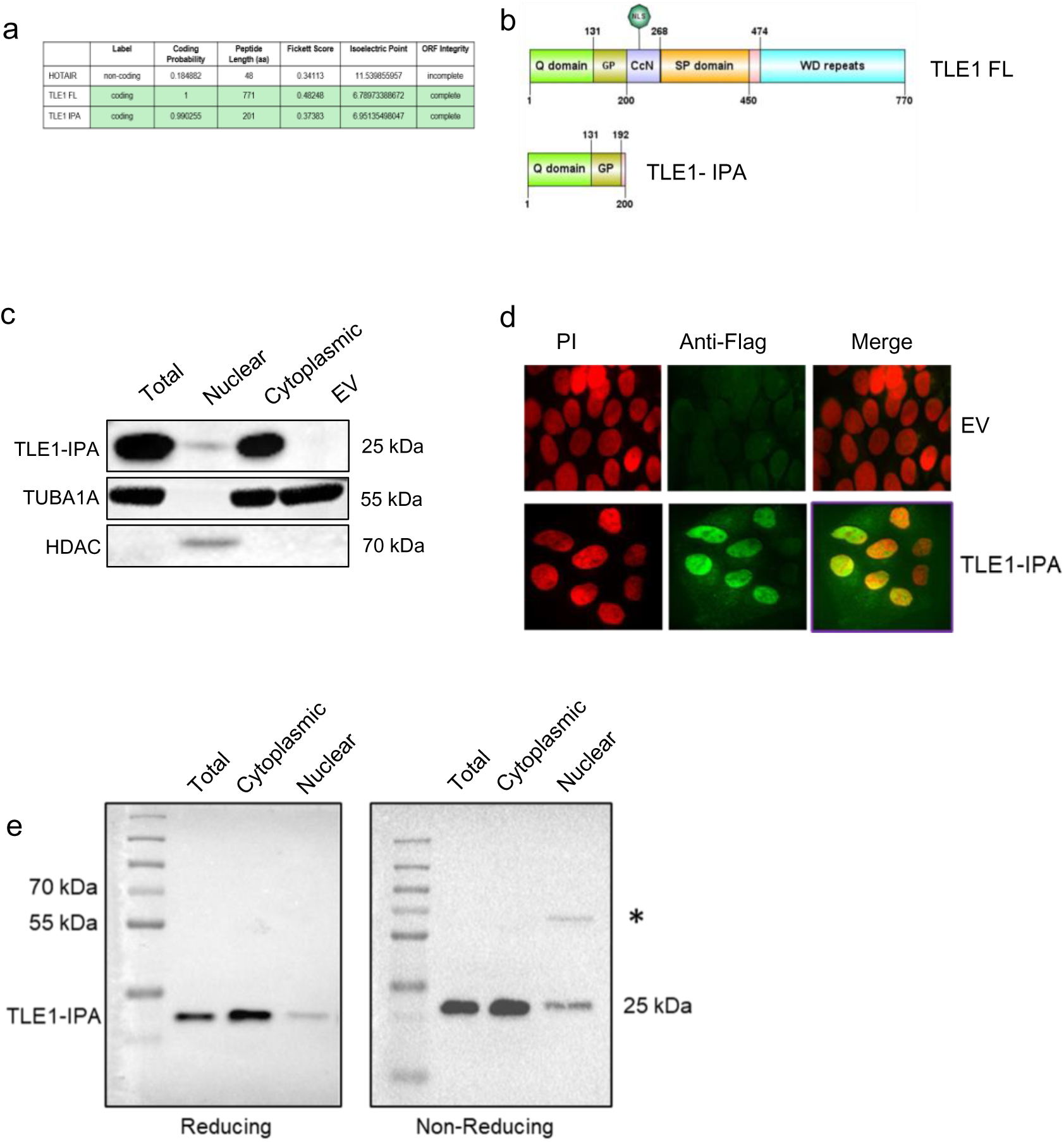
**a**. Coding potential of *TLE1*-IPA isoform is similar to *TLE1* FL. HOTAIR is a non-coding RNA, calculated by CPC2. **b**. Protein domains of FL and TLE1 IPA, **c**. Western blot showing expression of C-term truncated, N-Flag-TLE1-IPA in total, nuclear and cytoplasmic lysates of transfected MCF7 cells. The expected product size for TLE1-IPA is 25 kDa. TUBA1A (55 kDa) and HDAC (68 kDa) levels were used as cytoplasmic and nuclear protein controls. EV: empty pcDNA3.1 (-) vector, **d**. ICC for TLE1 IPA. Flag primary and Alexa Fluor 488 secondary antibodies were used to detect 3X-Flag-TLE1-IPA in transfected MCF7 cells. Propidium iodide was used for nuclear staining. Images were taken with ANDOR AMH200 DSD2 Spinning Disc microscope (100X) EV: empty vector-transfected cells.

Expanding on this observation, we aimed to confirm that the truncated TLE1 is translated. For this, we stably transfected MCF7 cells with an expression vector in which we cloned an N-terminal 3xFlag tag fused to the reading frame of the IPA isoform. The expression of approximately 25 kDa C-term truncated TLE1 protein was confirmed by western blotting (Figure 6c). We then performed immunocytochemistry (ICC), which confirmed the expression of truncated TLE1 (Figure 6d). Notably, despite lacking a nuclear localization signal (NLS), the truncated protein was detected in the nuclei of transfected cells (Figure 6c, d). Western blotting of subcellular fractions of lysates performed under non-reducing conditions confirmed not only the nuclear localization but also suggested the dimerization of tagged truncated proteins (Figure 6e). These results suggest that the truncated protein may dimerize with itself or other TLE proteins, facilitating its nuclear transport. To investigate the structural viability of this dimer, we used an enhanced AF2 sampling protocol through the ColabFold platform as described [23], [24], [25]. We used crystal structures of the human TLE1 Q-domain listed in the Protein Data Bank (PDB) (40M2 and 40M3) [39]. Identical to the crystal structure of the Q domain of full-length TLE1, AF2 predicted disordered regions in the first 20 and last 70 amino acids of the truncated protein, reflected by low pLDDT scores (Figure 7a, b). In contrast, the intact N-term Q-domain involving the dimerization region was modeled with high confidence. The Q domains in the full-length and truncated isoforms are identical; hence, binding between TLE proteins seems highly probable (Figure 7c). Indeed, the hydrophobic L26 and I29 residues at the N-term, required for dimerization/tetramerization, are also present in the truncated protein [39]. These results agreed with the non-reducing western blot results suggesting dimerization of Flag-tagged truncated TLE1 proteins (Figure 6e).

**Figure 7.**
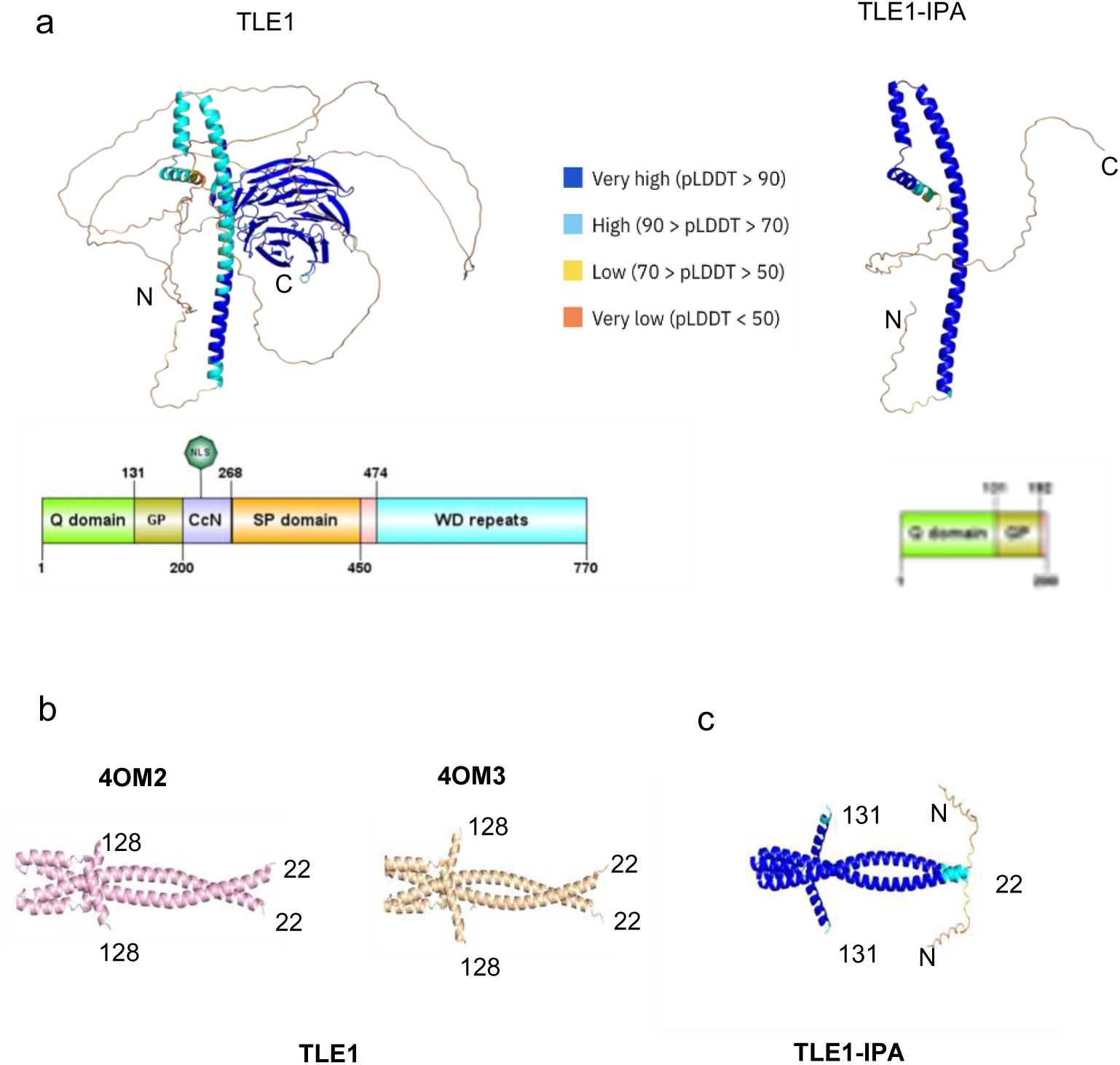
**a.**AF2 protein models of TLE1 and TLE1-IPA, colored according to AF2 plDDT confidence scores. Protein domain organizations are provided below the structural models. **b.** Available PDB structures of TLE1 dimers are shown. The left panel (pink) represents the A and C chains of 40M2 PDB, and the middle panel (wheat) shows the A and C chains of 40M3 PDB (Crystal structure of TLE1 N-terminal Q-domain dimers) **c**. The AF2 model of the TLE1 IPA dimers. plDDT confidence scores are provided in the key.

Consequently, the absence of C-terminal domains in the truncated protein is likely to alter interactions with other proteins, including pioneering factors for ERα. TLE proteins interact with many proteins with the WRPW/Y and Engrailed Homology 1 (EH1) (FxIxxI) motifs through their WDR interaction domain (2CE8, 2CE9) [40] (Figure 8a). Using massive sampling (AFsampling) to improve model quality for multimeric protein modeling [41], [42] we present the predicted interaction model of TLE1 WDR domain (light blue) and the EH1 motif of FOXG1 and FOXA1 (a pioneering factor for ERα) (Figure 8b). Based on these models, the truncated TLE1 protein cannot engage in WDR-dependent interactions, which may lead to alterations or disruptions in transcriptional regulatory networks. Of note, numerous known and predicted post-translational modification (PTM) sites (phosphorylation, acetylation, and ubiquitination) detected by PTMcode [43] reside between the 200-718 aa region (Table S3), which is also not present in the truncated protein.

**Figure 8.**
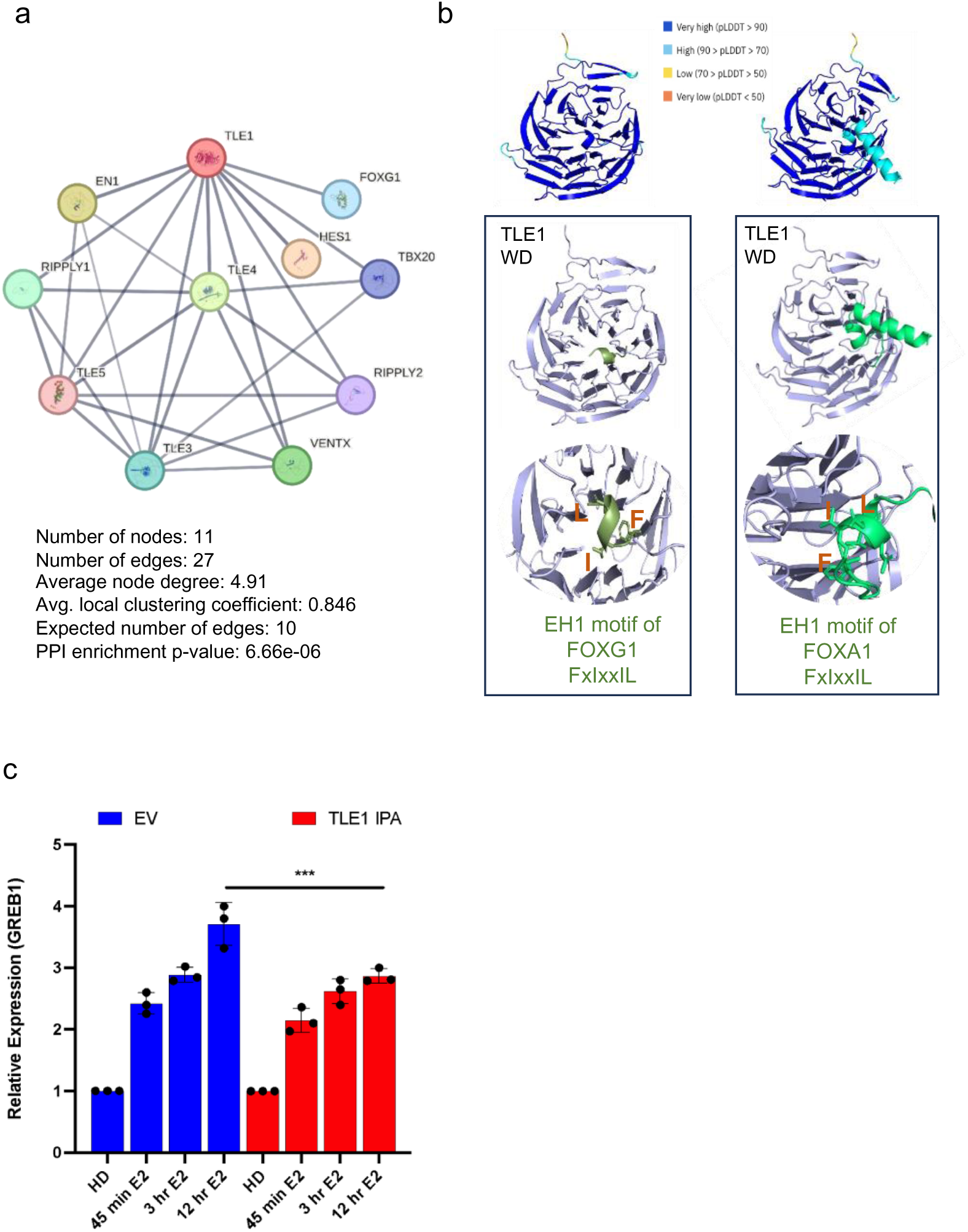
**a**. Protein-protein interaction network based on experimental evidence for TLE1. Figure created in STRING Database (STRING consortium 2024). FOXG1 is an interactor of TLE1. **b.** The left panel shows the predicted model of canonical TLE1 WDR domain and EH1 motif of FOXG1 (395-425) interaction by enhanced AF2 sampling. The confidence score of models is presented in color-coded pLDDT scores. The right panel shows the predicted model of the TLE1 WDR domain (light blue) and EH1 motif of FOXA1(lime green) (395-425), **c.** RT-qPCR showing E2-induced upregulation of *GREB1* in MCF7 cells that stably express TLE1-IPA compared with control cells (EV: Empty Vector). Both sets were hormone deprived (HD) for 72 hrs. and treated with E2 for the indicated time points. Expression of *GREB1* was normalized against the reference gene, *RPLP0.* GREB1 expression in HD cells was set to 1 (n=3 independent E2 treatments, one-way ANOVA, ns: non-significant, ****p<0.0001).

On the other hand, GREB1 has an N-terminal Zn-binding domain (Zn) followed by a circularly permuted catalytically inactive superfamily II (SFII) helicase domain and a C-terminal Fringe-like GT domain implicated in glycosylation (Figure S13). Through the GT domain, GREB1 catalyzes O-GlcNAcylation of ERα at residues T553/S554, which stabilizes ERα protein by inhibiting association with the ubiquitin ligase ZNF598. Hence, the E2 target gene GREB1 is in turn critical for ERα stabilization [44]. The experimentally confirmed *GREB1* IPA isoform we identified encodes for a C-term truncated GREB1 harboring the Zn domain but lacking the GT domain. However, AF2 could not predict high-confidence models for the truncated GREB1 (Figure S13), even though we could express an N-terminus FLAG-tagged truncated protein of GREB1 (Figure S13).

We also confirmed an additional IPA transcript that only resulted in a minor C-term truncation. An annotated IPA isoform (ENST00000343575) of *CXCL12* (C-X-C motif chemokine ligand 12) was upregulated in response to E2 stimulation in MCF7 cells. This isoform is 89 amino acids long compared to the 93 amino acids translated from the longer *CXCL12* isoform. The C-term truncation only involves 4 amino acids and no domain loss. Consequently, AF2 modeling did not predict a major change in protein structure (Figure S14).

Hence, based on the high-confidence protein models and significant domain loss, we focused on protein-level implications for the truncated TLE1.

### C-terminus truncated TLE1

TLE1 is known to enhance ERα-chromatin interactions, thereby promoting the binding of RNA polymerase II to a set of E2-responsive promoters, including *GREB1* [37]. To investigate the functionality of truncated TLE1, we analyzed the E2 response in MCF7 cells stably expressing the truncated protein (TLE1-IPA) compared to control cells transfected with an empty vector. MCF7 cells were first hormone-deprived for 72 hours and treated with either 100 nM E2 or ethanol (vehicle control) for 45 minutes, 3 and 12 hours, and RNA was isolated. Next, we evaluated the upregulation of *GREB1* by RT-qPCR. Earlier *GREB1* was identified as a TLE1-dependent and E2-responsive gene [37]. Notably, the upregulation of *GREB1* in cells overexpressing the truncated protein was lower than in control cells that did not overexpress the truncated TLE1 (Figure 8c). These results suggest that the truncated TLE1 may impair the TLE-dependent E2 response, including *GREB1* transcription. This impairment is likely caused by disruptions in critical protein interactions, as predicted by AF2.

These findings prompt new questions regarding the functional role of the truncated TLE1 and other protein isoforms in both physiological and pathological contexts. Elucidating the mechanism of action and protein interactions of truncated TLE1 could be important in breast cancer progression and endocrine therapy response. These insights may apply to other transcriptional networks regulated by TLE1 and to other truncated proteins translated from IPA isoforms.

## Discussion

Long-read sequencing offers the significant advantage of generating reads that typically encompass the entire length of most transcripts [45], allowing the sequencing of complete isoforms in continuous reads. However, it also presents challenges, including high sequencing error rates and smaller library sizes, which can reduce the quantification accuracy. In parallel, several specialized long-read tools have been developed for high-confidence isoform calling. FLAIR, for instance, is a reference-free method that maps the reads to the reference genome, clusters alignments into groups, and collapses them into isoforms [12]. Other computational tools are also designed to improve the accuracy of isoform calling [46]. Given the challenges associated with long-read sequencing and the accuracy of computational tools, we opted to complement and validate our findings using additional approaches, including 3’-seq and experimental validation. The dynamic regulation patterns of gene expression observed in our study align with the well-established E2-regulated gene expression profiles. Furthermore, our findings emphasize previously unrecognized aspects of transcriptome complexity, shedding new light on the intricate mechanisms involved.

Looking deeper into the transcriptomic complexity, we identified non-coding RNAs (ncRNAs) regulated by E2, predominantly long non-coding RNAs (lncRNAs). Interestingly, while most ncRNAs were initially upregulated, a notable increase in downregulated non-coding transcripts was observed after 12 hours. For example, our combinatorial approach revealed significant downregulation of lncGATA3-7 upon E2 treatment, which has a breast cancer subtype expression. The enrichment processes such as apoptosis and gland development underscore the potential roles of ncRNAs in cancer biology, particularly in the context of ER+ breast cancer.

For protein-coding genes, we identified a complex landscape of differentially expressed alternative splicing isoforms due to exon splicing or intron retention. These alternative isoforms could have significant implications at the protein level, related to their functions or with implications for their detection. Earlier we identified an alternatively spliced PD-L1 isoform, PD-L1Δ3 which is expressed in breast tumors but cannot interact with PD-1. However, PD-L1Δ3 is not detectable by most of the PD-L1 immunohistochemical assays or antibodies currently used in clinical practice. As a result, its presence potentially affects treatment decisions and patient outcomes in immunotherapy [47]. These findings underscore the importance of uncovering transcript isoforms, expanding our understanding of their impact on protein function and therapeutic interventions.

As for isoforms with alternate 3’-ends, global alternative polyadenylation and variations in 3′UTR lengths were initially observed in activated T cells and cancer cells [48], [49]. But beyond 3’UTRs, approximately 20% of human genes have at least one intronic polyadenylation event that can potentially lead to mRNA isoforms [50]. Consequently, intronic polyadenylation profiles in the pan-cancer transcriptome are also beginning to emerge [51], [52]. When we looked at transcriptome diversity from a 3’-end perspective, we were surprised to identify numerous intronically polyadenylated transcripts in E2-treated cells. These transcripts either terminate in early or mid-intronic regions. A notable number of EI transcripts detected by DRS, originate from genes with long introns. Functional roles and/or implications for these transcripts and their effect on full-length isoforms remain to be investigated. The 3’-seq reads that were not captured by DRS also originated from longer introns that were enriched for Alu elements and genomic A stretch regions. Internal priming during library preparation may have enhanced the detection of these regions and these reads may have originated from pre-mRNAs due to delayed splicing events and potentially modulating the full-length mRNA levels.

For IPA transcripts that are likely to generate truncated proteins, earlier examples were reported for dominant-negative and secreted variants of receptor tyrosine kinases [53]. Since then, subsequent examples of truncated protein have been known [54], [55]. In fact, in cancers, most truncated proteins are not caused by genetic mutations but instead, result from intronic polyadenylation, and they typically exist as minor isoforms in normal tissues. Previously, we identified a C-terminally truncated EIF2Bγ protein produced by a minor isoform using a different approach. This truncated protein disrupts the EIF2Bγ-EIF2γ interaction and could be crucial for regulating stress-induced translation in both normal and disease states [56]. Such findings underscore the broader significance of intronically terminated transcripts, which can serve diverse functions [57].

Building on this understanding, we concentrated on the MI group of E2-regulated transcripts, specifically focusing on the *GREB1* and *TLE1* IPA isoforms. Both genes are integral to the E2 response, making their alternative isoforms valuable targets for exploring the nuances of transcriptional regulation and functional outcomes in the E2-regulated transcriptome. We confirmed the intronic termination sites for both IPA isoforms and detected comparable half-lives to their full-length isoforms. The expression of both IPA isoforms was detected in single-cell RNA sequences. For TLE1, the DRS captured IPA isoform’s 3’-end was identical to a known transcript, TLE1-201, which was not assigned a continuous reading frame because it lacked sequences from the 5’-end. Because we got the complete sequence from DRS, we regarded the DRS-captured IPA isoform as the complete version of TLE1-201 and showed that it co-exists with the full-length isoform in the TCGA breast cancer cohort. Curiously, lower IPA/FL isoform levels are associated with worse survival in ER+ breast cancer patients, indicating functional relevance. The structural models predicted by AF2 showed that the truncated TLE1 protein retains the essential Q domain, crucial for dimerization, but lacked other C-terminal domains, potentially altering its interaction capabilities with key regulatory proteins, including FOXG1 and FOXA1. Supporting this, we experimentally showed dimerization and nuclear localization of the Flag-tagged truncated TLE1, which also led to decreased upregulation of *GREB1* mRNA in response to E2 stimulation. Hence, the absence of the WDR interaction domain in the truncated protein seems to cause a complete loss or weakening of protein interactions, disrupting the transcriptional network. Further work on the truncated TLE1 will reveal the extent of its effects on gene expression in normal and disease states. For the C-term truncated GREB1, AF2 did not predict high-confidence models but the loss of glycosylation domain is likely to have consequences.

In summary, our findings reveal new complexities in the E2-regulated transcriptome, particularly IPAs and truncated proteins, advancing our understanding of gene regulation. Growing evidence highlights the functional significance of transcript isoforms, which can alter the activity of their full-length counterparts. Despite their importance, these intronically isoforms are often overlooked due to detection challenges. Long-read sequencing and emerging analysis methods now enable more accurate de novo assembly, enhanced mapping precision, and better isoform identification, allowing deeper exploration of previously unexplored protein-level implications. These advancements offer potential therapeutic implications by uncovering insights into gene regulation that were once difficult to address.

## Data Availability

Sequencing data generated for this study are available at NCBI’s Sequence Read Archive through the BioProject accession numbers PRJNA1195033 and PRJNA1196847.

## Author contributions

DND performed bioinformatic analysis for ONT and 3’seq, IO performed ONT sequencing, contributed to bioinformatic analysis, performed experiments, IY performed computational structure analysis and experiments, EA, IE, UCY, DK, and EC performed experiments, MC prepared 3’seq libraries, EK and TC supervised in silico work, AEEB conceptualized the work, drafted and reviewed the manuscript. All authors contributed to the writing of the manuscript.

## Acknowledgments

This project was funded by TUBITAK 120Z464. This article is based upon work from COST Action CA21154 TRANSLACORE, supported by COST (European Cooperation in Science and Technology). EK and IY are supported by the EMBO Installation Grant (No: 4421). EA was supported by TUBITAK BIDEB 2210A. We thank Dr. Ekim Taskiran (Hacettepe University) for 3’-sequencing and Dr. Cagdas Devrim Son (METU) for his help in microscopy. We thank Busra Savas for her assistance in enhanced AF2 sampling and Dr. Oguzhan Begik for insightful discussions on nanopore DRS sequencing.

## References

[1] N. Hah and W. L. Kraus, “Hormone-regulated transcriptomes: lessons learned from estrogen signaling pathways in breast cancer cells.,” Mol Cell Endocrinol, vol. 382, no. 1, pp. 652–664, Jan. 2014, doi: 10.1016/j.mce.2013.06.021.

[2] N. Hah et al., “A rapid, extensive, and transient transcriptional response to estrogen signaling in breast cancer cells.,” Cell, vol. 145, no. 4, pp. 622–634, May 2011, doi: 10.1016/j.cell.2011.03.042.

[3] Z. Li et al., “The EstroGene Database Reveals Diverse Temporal, Context-Dependent, and Bidirectional Estrogen Receptor Regulomes in Breast Cancer.,” Cancer Res, vol. 83, no. 16, pp. 2656–2674, Aug. 2023, doi: 10.1158/0008-5472.CAN-23-0539.

[4] A. Kalsotra and T. A. Cooper, “Functional consequences of developmentally regulated alternative splicing.,” Nat Rev Genet, vol. 12, no. 10, pp. 715–729, Sep. 2011, doi: 10.1038/nrg3052.

[5] S. A. Bustin et al., “The MIQE guidelines: minimum information for publication of quantitative real-time PCR experiments.,” Clin Chem, vol. 55, no. 4, pp. 611–622, Apr. 2009, doi: 10.1373/clinchem.2008.112797.

[6] M. Erdem et al., “Identification of an mRNA isoform switch for HNRNPA1 in breast cancers.,” Sci Rep, vol. 11, no. 1, p. 24444, Dec. 2021, doi: 10.1038/s41598-021-04007-y.

[7] A. R. Quinlan and I. M. Hall, “BEDTools: a flexible suite of utilities for comparing genomic features.,” Bioinformatics, vol. 26, no. 6, pp. 841–842, Mar. 2010, doi: 10.1093/bioinformatics/btq033.

[8] Y. Zhang et al., “Model-based analysis of ChIP-Seq (MACS).,” Genome Biol, vol. 9, no. 9, p. R137, 2008, doi: 10.1186/gb-2008-9-9-r137.

[9] E. Beaudoing, S. Freier, J. R. Wyatt, J. M. Claverie, and D. Gautheret, “Patterns of variant polyadenylation signal usage in human genes.,” Genome Res, vol. 10, no. 7, pp. 1001–1010, Jul. 2000, doi: 10.1101/gr.10.7.1001.

[10] N. J. Proudfoot, “Ending the message: poly(A) signals then and now.,” Genes Dev, vol. 25, no. 17, pp. 1770–1782, Sep. 2011, doi: 10.1101/gad.17268411.

[11] C. J. Herrmann, R. Schmidt, A. Kanitz, P. Artimo, A. J. Gruber, and M. Zavolan, “PolyASite 2.0: a consolidated atlas of polyadenylation sites from 3’ end sequencing.,” Nucleic Acids Res, vol. 48, no. D1, pp. D174–D179, Jan. 2020, doi: 10.1093/nar/gkz918.

[12] A. D. Tang et al., “Full-length transcript characterization of SF3B1 mutation in chronic lymphocytic leukemia reveals downregulation of retained introns.,” Nat Commun, vol. 11, no. 1, p. 1438, Mar. 2020, doi: 10.1038/s41467-020-15171-6.

[13] S. Zhang et al., “RNAenrich: a web server for non-coding RNA enrichment.,” Bioinformatics, vol. 39, no. 7, Jul. 2023, doi: 10.1093/bioinformatics/btad421.

[14] A. Subramanian et al., “Gene set enrichment analysis: a knowledge-based approach for interpreting genome-wide expression profiles.,” Proc Natl Acad Sci U S A, vol. 102, no. 43, pp. 15545–15550, Oct. 2005, doi: 10.1073/pnas.0506580102.

[15] S. Fleige, V. Walf, S. Huch, C. Prgomet, J. Sehm, and M. W. Pfaffl, “Comparison of relative mRNA quantification models and the impact of RNA integrity in quantitative real-time RT-PCR.,” Biotechnol Lett, vol. 28, no. 19, pp. 1601–1613, Oct. 2006, doi: 10.1007/s10529-006-9127-2.

[16] W. Chung et al., “Single-cell RNA-seq enables comprehensive tumour and immune cell profiling in primary breast cancer.,” Nat Commun, vol. 8, p. 15081, May 2017, doi: 10.1038/ncomms15081.

[17] W. C. Reinhold, M. Sunshine, S. Varma, J. H. Doroshow, and Y. Pommier, “Using CellMiner 1.6 for Systems Pharmacology and Genomic Analysis of the NCI-60.,” Clin Cancer Res, vol. 21, no. 17, pp. 3841–3852, Sep. 2015, doi: 10.1158/1078-0432.CCR-15-0335.

[18] J. Baran-Gale, J. E. Purvis, and P. Sethupathy, “An integrative transcriptomics approach identifies miR-503 as a candidate master regulator of the estrogen response in MCF-7 breast cancer cells.,” RNA, vol. 22, no. 10, pp. 1592–1603, Oct. 2016, doi: 10.1261/rna.056895.116.

[19] S. Al Saleh, F. Al Mulla, and Y. A. Luqmani, “Estrogen receptor silencing induces epithelial to mesenchymal transition in human breast cancer cells.,” PLoS One, vol. 6, no. 6, p. e20610, 2011, doi: 10.1371/journal.pone.0020610.

[20] Y.-J. Kang et al., “CPC2: a fast and accurate coding potential calculator based on sequence intrinsic features.,” Nucleic Acids Res, vol. 45, no. W1, pp. W12–W16, Jul. 2017, doi: 10.1093/nar/gkx428.

[21] “UniProt: the Universal Protein Knowledgebase in 2023.,” Nucleic Acids Res, vol. 51, no. D1, pp. D523–D531, Jan. 2023,doi: 10.1093/nar/gkac1052.

[22] H. M. Berman et al., “The Protein Data Bank.,” Nucleic Acids Res, vol. 28, no. 1, pp. 235– 242, Jan. 2000, doi: 10.1093/nar/28.1.235.

[23] J. Jumper et al., “Highly accurate protein structure prediction with AlphaFold.,” Nature, vol. 596, no. 7873, pp. 583–589, Aug. 2021, doi: 10.1038/s41586-021-03819-2.

[24] M. Mirdita, K. Schütze, Y. Moriwaki, L. Heo, S. Ovchinnikov, and M. Steinegger, “ColabFold: making protein folding accessible to all,” Nat Methods, vol. 19, no. 6, Art. no. 6, Jun. 2022, doi: 10.1038/s41592-022-01488-1.

[25] B. Savaş, İ. Yılmazbilek, A. Özsan, and E. Karaca, “Towards a greener AlphaFold2 protocol for protein complex modeling: Insights from CAPRI Round 55,” Oct. 11, 2024, *bioRxiv*. doi: 10.1101/2024.10.07.616947.

[26] Y. Xie et al., “IBS 2.0: an upgraded illustrator for the visualization of biological sequences.,” Nucleic Acids Res, vol. 50, no. W1, pp. W420–W426, Jul. 2022, doi: 10.1093/nar/gkac373.

[27] A. Prokesch and M. A. Lazar, “A hormone sends instant messages to the genome.,” Cell, vol. 145, no. 4, pp. 499–501, May 2011, doi: 10.1016/j.cell.2011.04.016.

[28] K. Basu, A. Dey, and M. Kiran, “Inefficient splicing of long non-coding RNAs is associated with higher transcript complexity in human and mouse,” RNA Biol, vol. 20, no. 1, pp. 563–572, Jan. 2023, doi: 10.1080/15476286.2023.2242649.

[29] S. J. Liu et al., “Single-cell analysis of long non-coding RNAs in the developing human neocortex.,” Genome Biol, vol. 17, p. 67, Apr. 2016, doi: 10.1186/s13059-016-0932-1.

[30] W. Sun et al., “Transcriptome analysis of luminal breast cancer reveals a role for LOL in tumor progression and tamoxifen resistance.,” Int J Cancer, vol. 145, no. 3, pp. 842–856, Aug. 2019, doi: 10.1002/ijc.32185.

[31] J. S. Mattick et al., “Long non-coding RNAs: definitions, functions, challenges and recommendations.,” Nat Rev Mol Cell Biol, vol. 24, no. 6, pp. 430–447, Jun. 2023, doi: 10.1038/s41580-022-00566-8.

[32] P. Li et al., “CRB3 downregulation confers breast cancer stem cell traits through TAZ/β-catenin.,” Oncogenesis, vol. 6, no. 4, p. e322, Apr. 2017, doi: 10.1038/oncsis.2017.24.

[33] P. A. Cordell, T. S. Futers, P. J. Grant, and R. J. Pease, “The Human hydroxyacylglutathione hydrolase (HAGH) gene encodes both cytosolic and mitochondrial forms of glyoxalase II.,” J Biol Chem, vol. 279, no. 27, pp. 28653–28661, Jul. 2004, doi: 10.1074/jbc.M403470200.

[34] P. Heyn, A. T. Kalinka, P. Tomancak, and K. M. Neugebauer, “Introns and gene expression: cellular constraints, transcriptional regulation, and evolutionary consequences.,” Bioessays, vol. 37, no. 2, pp. 148–154, Feb. 2015, doi: 10.1002/bies.201400138.

[35] S. Lee et al., “Covering all your bases: incorporating intron signal from RNA-seq data.,” NAR Genom Bioinform, vol. 2, no. 3, p. lqaa073, Sep. 2020, doi: 10.1093/nargab/lqaa073.

[36] J. M. Rae, M. D. Johnson, J. O. Scheys, K. E. Cordero, J. M. Larios, and M. E. Lippman, “GREB 1 is a critical regulator of hormone dependent breast cancer growth.,” Breast Cancer Res Treat, vol. 92, no. 2, pp. 141–149, Jul. 2005, doi: 10.1007/s10549-005-1483-4.

[37] K. A. Holmes et al., “Transducin-like enhancer protein 1 mediates estrogen receptor binding and transcriptional activity in breast cancer cells.,” Proc Natl Acad Sci U S A, vol. 109, no. 8, pp. 2748–2753, Feb. 2012, doi: 10.1073/pnas.1018863108.

[38] D. J. Herrick and J. Ross, “The half-life of c-myc mRNA in growing and serum-stimulated cells: influence of the coding and 3’ untranslated regions and role of ribosome translocation.,” Mol Cell Biol, vol. 14, no. 3, pp. 2119–2128, Mar. 1994, doi: 10.1128/mcb.14.3.2119-2128.1994.

[39] J. V. Chodaparambil et al., “Molecular functions of the TLE tetramerization domain in Wnt target gene repression.,” EMBO J, vol. 33, no. 7, pp. 719–731, Apr. 2014, doi: 10.1002/embj.201387188.

[40] K. E. Nye, K. A. Knox, and A. J. Pinching, “Lymphocytes from HIV-infected individuals show aberrant inositol polyphosphate metabolism which reverses after zidovudine therapy.,” AIDS, vol. 5, no. 4, pp. 413–417, Apr. 1991, doi: 10.1097/00002030-199104000-00009.

[41] A. Elofsson, “Progress at protein structure prediction, as seen in CASP15.,” Curr Opin Struct Biol, vol. 80, p. 102594, Jun. 2023, doi: 10.1016/j.sbi.2023.102594.

[42] B. Wallner, “AFsample: improving multimer prediction with AlphaFold using massive sampling.,” Bioinformatics, vol. 39, no. 9, Sep. 2023, doi: 10.1093/bioinformatics/btad573.

[43] P. Minguez, I. Letunic, L. Parca, and P. Bork, “PTMcode: a database of known and predicted functional associations between post-translational modifications in proteins.,” Nucleic Acids Res, vol. 41, no. Database issue, pp. D306-311, Jan. 2013, doi: 10.1093/nar/gks1230.

[44] E. M. Shin et al., “GREB1: An evolutionarily conserved protein with a glycosyltransferase domain links ERα glycosylation and stability to cancer,” Sci Adv, vol. 7, no. 12, p. eabe2470, Mar. 2021, doi: 10.1126/sciadv.abe2470.

[45] X. Dong et al., “The long and the short of it: unlocking nanopore long-read RNA sequencing data with short-read differential expression analysis tools.,” NAR Genom Bioinform, vol. 3, no. 2, p. lqab028, Jun. 2021, doi: 10.1093/nargab/lqab028.

[46] Y. Su et al., “Comprehensive assessment of mRNA isoform detection methods for long-read sequencing data.,” Nat Commun, vol. 15, no. 1, p. 3972, May 2024, doi: 10.1038/s41467-024-48117-3.

[47] D. N. Dioken, I. Ozgul, I. Yilmazbilek, M. C. Yakicier, E. Karaca, and A. E. Erson-Bensan, “An alternatively spliced PD-L1 isoform PD-L1Δ3, and PD-L2 expression in breast cancers: implications for eligibility scoring and immunotherapy response.,” Cancer Immunol Immunother, vol. 72, no. 12, pp. 4065–4075, Dec. 2023, doi: 10.1007/s00262-023-03543-y.

[48] C. Mayr and D. P. Bartel, “Widespread shortening of 3’UTRs by alternative cleavage and polyadenylation activates oncogenes in cancer cells,” Cell, vol. 138, no. 4, pp. 673–684, Aug. 2009, doi: 10.1016/j.cell.2009.06.016.

[49] R. Sandberg, J. R. Neilson, A. Sarma, P. A. Sharp, and C. B. Burge, “Proliferating cells express mRNAs with shortened 3’ untranslated regions and fewer microRNA target sites.,” Science, vol. 320, no. 5883, pp. 1643–1647, Jun. 2008, doi: 10.1126/science.1155390.

[50] B. Tian, Z. Pan, and J. Y. Lee, “Widespread mRNA polyadenylation events in introns indicate dynamic interplay between polyadenylation and splicing.,” Genome Res, vol. 17, no. 2, pp. 156–165, Feb. 2007, doi: 10.1101/gr.5532707.

[51] O. Begik, M. Oyken, T. Cinkilli Alican, T. Can, and A. E. Erson-Bensan, “Alternative Polyadenylation Patterns for Novel Gene Discovery and Classification in Cancer,” Neoplasia, vol. 19, no. 7, pp. 574–582, Jul. 2017, doi: 10.1016/j.neo.2017.04.008.

[52] J. Sun et al., “Dichotomous intronic polyadenylation profiles reveal multifaceted gene functions in the pan-cancer transcriptome.,” Exp Mol Med, vol. 56, no. 10, pp. 2145–2161, Oct. 2024, doi: 10.1038/s12276-024-01289-w.

[53] S. Vorlová et al., “Induction of antagonistic soluble decoy receptor tyrosine kinases by intronic polyA activation.,” Mol Cell, vol. 43, no. 6, pp. 927–939, Sep. 2011, doi: 10.1016/j.molcel.2011.08.009.

[54] A. E. Erson-Bensan and T. Can, “Alternative Polyadenylation: Another Foe in Cancer.,” Mol Cancer Res, vol. 14, no. 6, pp. 507–517, Jun. 2016, doi: 10.1158/1541-7786.MCR-15-0489.

[55] N. K. Mohanan, F. Shaji, G. R. Koshre, and R. S. Laishram, “Alternative polyadenylation: An enigma of transcript length variation in health and disease,” Wiley Interdiscip Rev RNA, vol. 13, no. 1, p. e1692, Jan. 2022, doi: 10.1002/wrna.1692.

[56] A. Circir et al., “A C-term truncated EIF2Bγ protein encoded by an intronically polyadenylated isoform introduces unfavorable EIF2Bγ-EIF2γ interactions.,” Proteins, vol. 90, no. 3, pp. 889–897, Mar. 2022, doi: 10.1002/prot.26284.

[57] K. Kamieniarz-Gdula and N. J. Proudfoot, “Transcriptional Control by Premature Termination: A Forgotten Mechanism.,” Trends Genet, vol. 35, no. 8, pp. 553–564, Aug. 2019, doi: 10.1016/j.tig.2019.05.005.

