## Supplementary material for "E2-Regulated Transcriptome Complexity Revealed by Long-Read Direct RNA Sequencing: From Isoform Discovery to Truncated Proteins": Supp. Fig

### Supplementary Figures

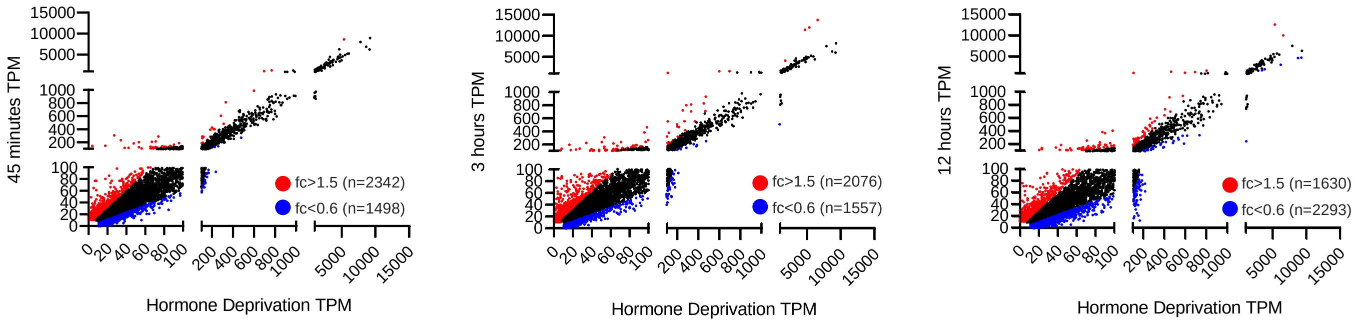


**Figure S1.** DRS detected differential gene expression (TPM) upon E2 stimulation for 45 minutes, 3, and 12 hours in MCF7 cells compared with hormone-deprived control cells. Red dots indicate significantly upregulated genes (fold change>1.5), blue dots represent significantly downregulated genes (fold change < 0.6), and black dots denote genes with fold changes between these thresholds.

**a**

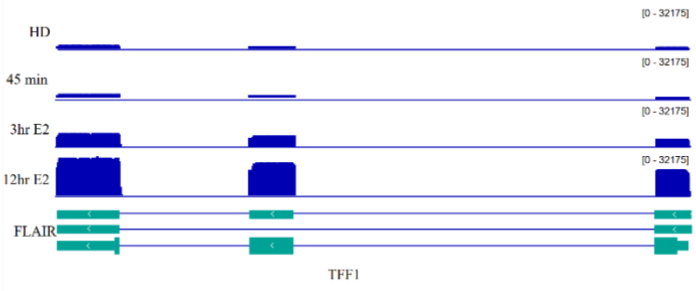


**b**

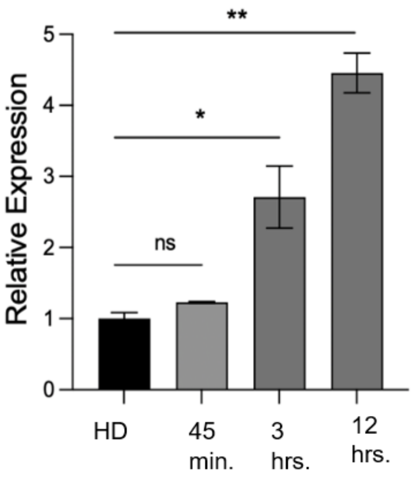


**Figure S2.** *TFF1* upregulation in MCF7 cells upon E2 treatment.

**a.** IGV illustration of *TFF1* DRS reads and FLAIR identified transcripts across different time points of E2 treatment, **b**. RT-qPCR confirmation of *TFF1* upregulation in response to E2. *RPLP0* was used as the reference gene. HD: Hormone-deprived control cells. Statistical significance was evaluated by ANOVA, with * $p < 0.05$, ** $p < 0.01$, and ns: not significant.

a

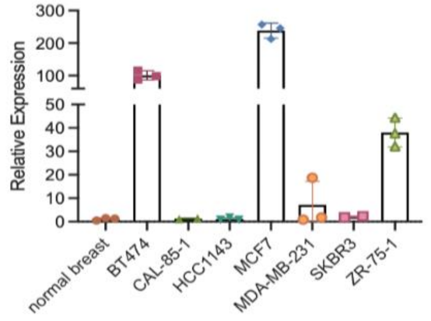


| Cell lines | ER | PR | HER2 |
|---|---|---|---|
| MCF7 | + | + | - |
| ZR751 | + | +/- | - |
| BT474 | + | + | + |
| SKBR3 | - | - | + |
| HCC1143 | - | - | - |
| CAL851 | - | - | - |
| MDAMB231 | - | - | - |

b

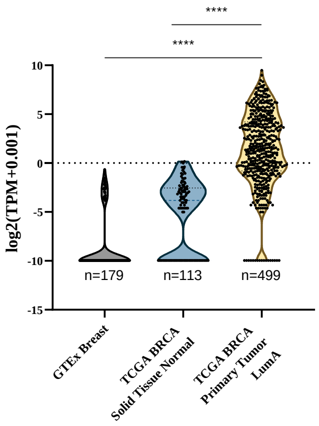


c

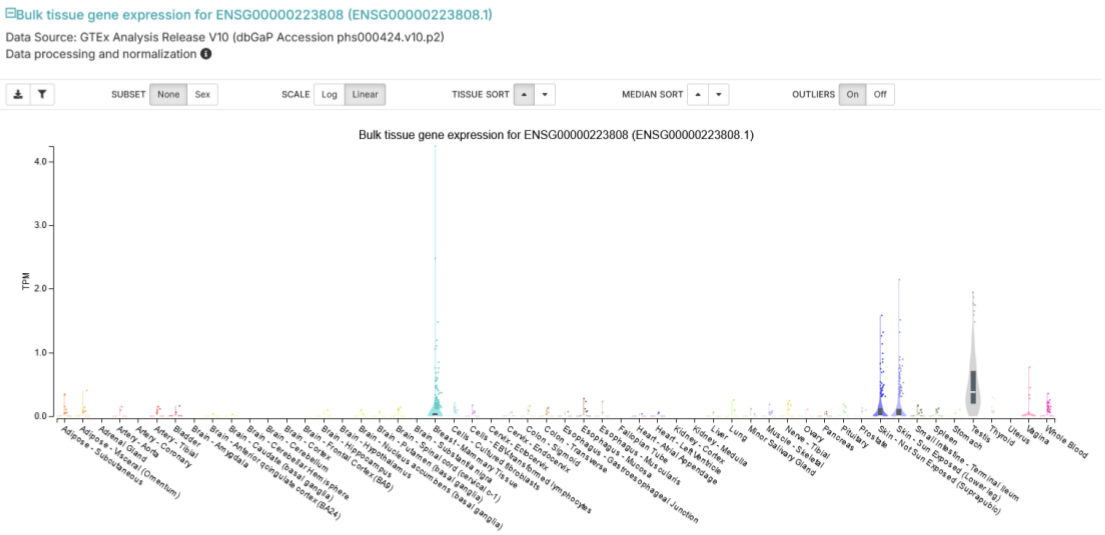


**Figure S3.** LncGATA3-7 expression

**a**. *LncGATA3-7* expression in breast cancer cell lines (MDA-MB-231, HCC1143, CAL-85-1, SK-BR-3, BT474, MCF7, ZR-75-1). *RPLP0* expression was used for normalization. **b.** Expression levels of lncGATA3-7 across three datasets: GTEx Breast (Normal Tissue), TCGA BRCA Solid Tissue Normal, and TCGA BRCA “Luminal A” Primary Tumors. GTEx Breast represents healthy breast tissue. TCGA BRCA Solid Tissue Normal includes non-tumorous breast tissue adjacent to tumors. TCGA BRCA “Luminal A” Primary Tumor corresponds to primary breast cancer samples classified under the Luminal A subtype, **c.** GTEx tissue expression of ENSG00000223808 (https://gtexportal.org/) (One way ANOVA, multiple comparisons, ****p<0.0001.

**a**

**b**

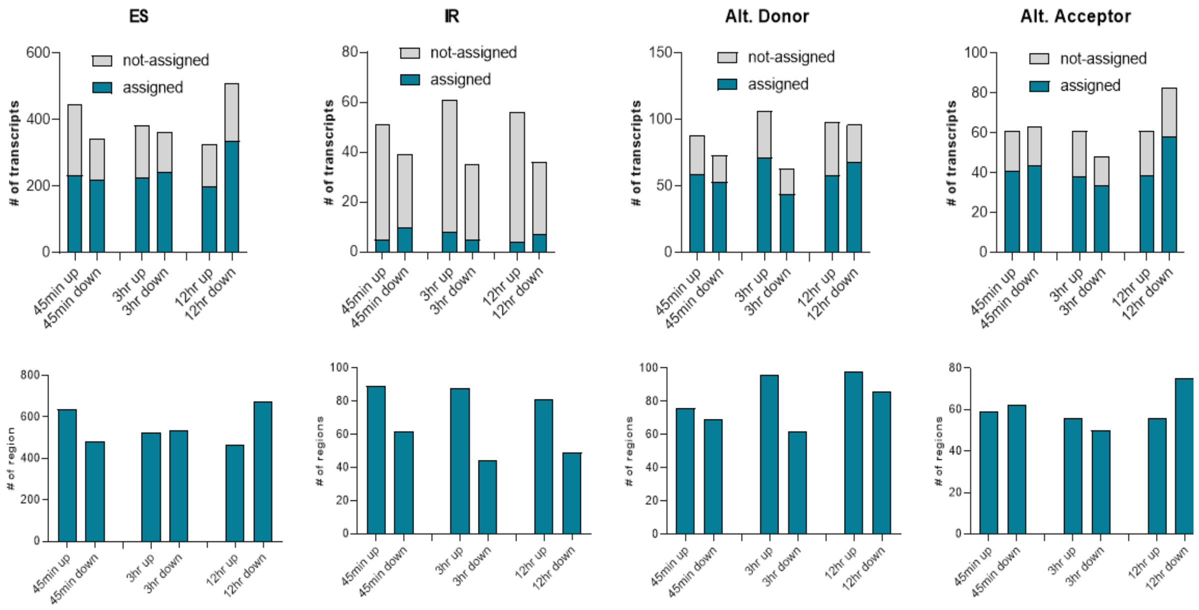


**Figure S4.** Splicing events in differentially expressed transcripts across all time points.

**a.** The number of differentially expressed transcripts (fold change >1.5 or fold change <0.6) containing specific splicing events at each time point of E2 induction. Transcripts are filtered to include only those with TPM ≥ 10 at all time points and with a ratio ≥ 10% relative to other transcripts of the same gene. Annotated transcripts are depicted as blue bars, and not-annotated transcripts are depicted as grey bars. **b.** The number of genomic regions implicated in splicing events of differentially expressed genes (fold change >1.5 or <0.6) at each time point of E2 treatment. Transcripts are filtered to include only those with TPM ≥ 10 at all-time points and with a ratio ≥ 10% relative to other transcripts of the same gene.

a

b

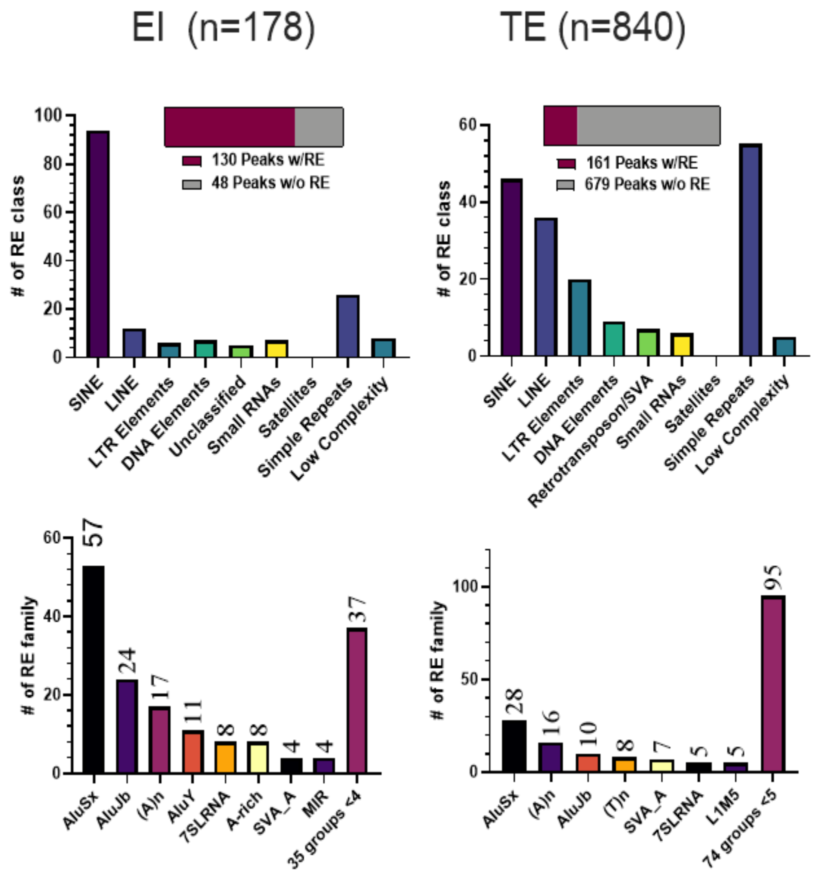


**Figure S5.** 3'seq reads originating from early intronic (EI) regions. **a.** Repeat elements (RE) located ± 100 bp peak ends of EI (EI, n = 178) and Terminal Exon (TE, n=840) transcripts, determined by RepeatMaster (https://www.repeatmasker.org). The distribution bar depicts the number of peaks (reads) with (burgundy) or without REs (gray) across all time points, **b**. Number and distribution of RE families in EI (left) and TE (right) genes.

**a**

TPD52L1

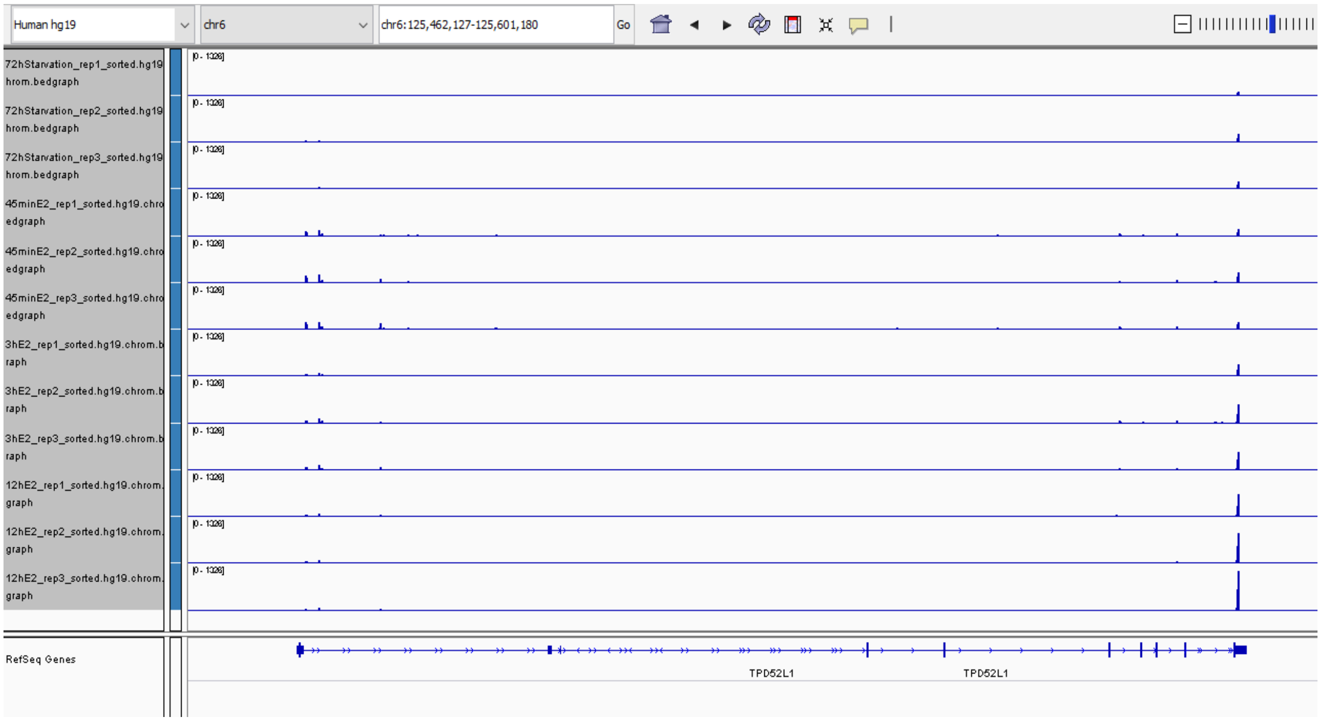


IGFBP4

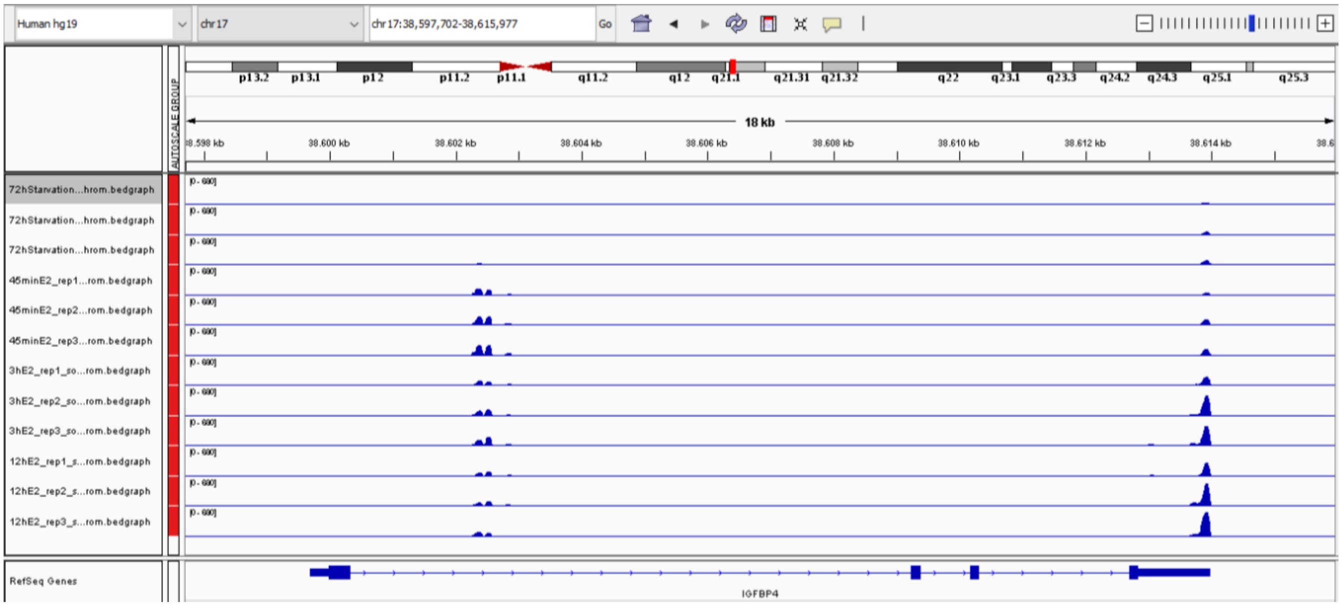

**b**

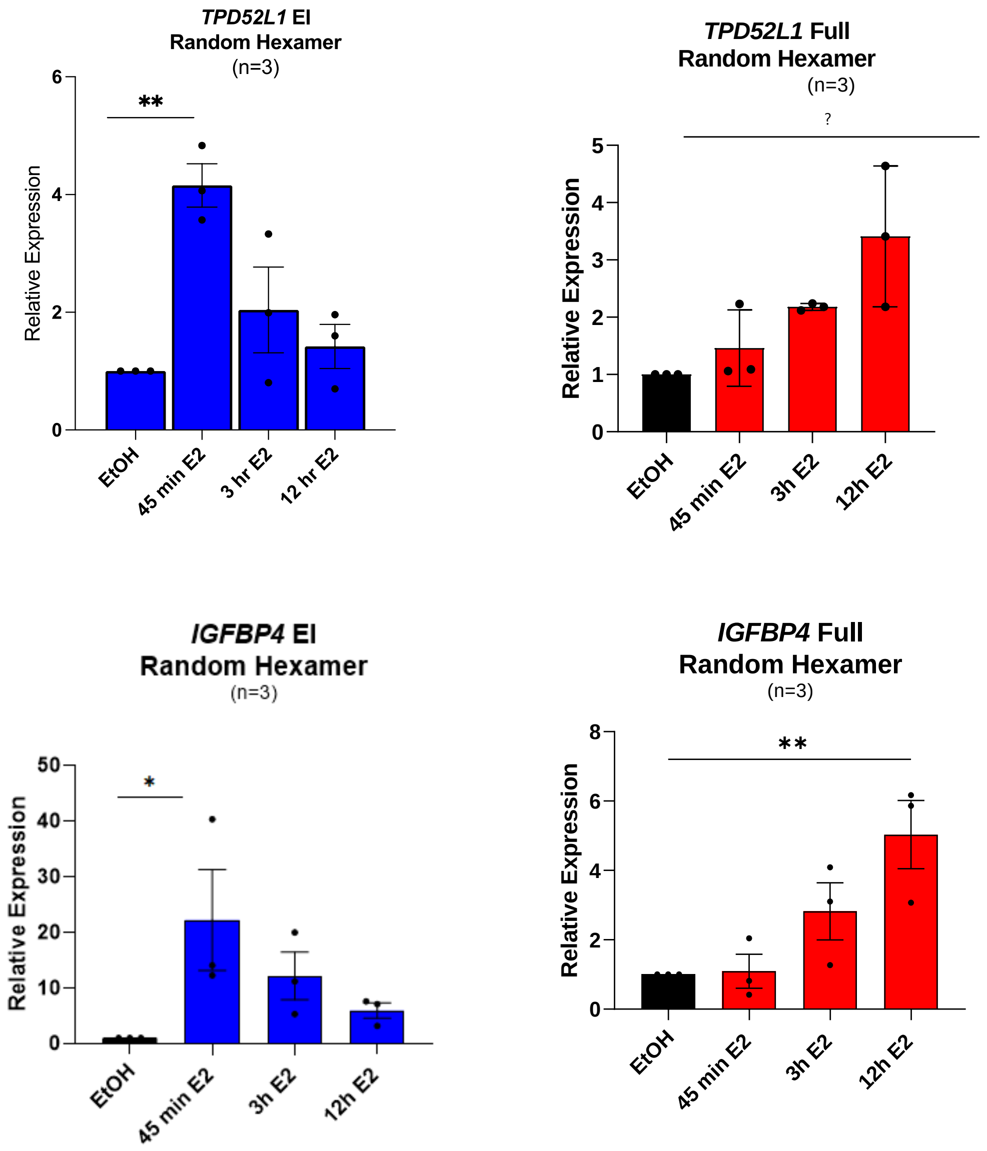


**Figure S6. a**. IGV display of 3'seq reads from *TPD52L1* and *IGFBP4* early intron and terminal exon regions, **b**. RT-qPCR results for EI (early intron) region and full-length using random-hexamer primed cDNA templates. Cells were hormone-deprived for 72 hours and treated with E2 for indicated durations. Expression from each time point was normalized to the vehicle control (EtOH: ethanol) samples of respective time points. *RLPL0* levels were used for normalization purposes (n=3 independent E2 treatments, one-way ANOVA, *p<0.05, **p<0.001).

a

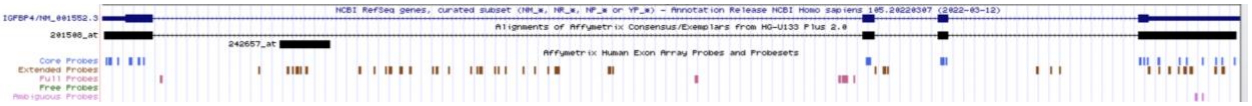


b

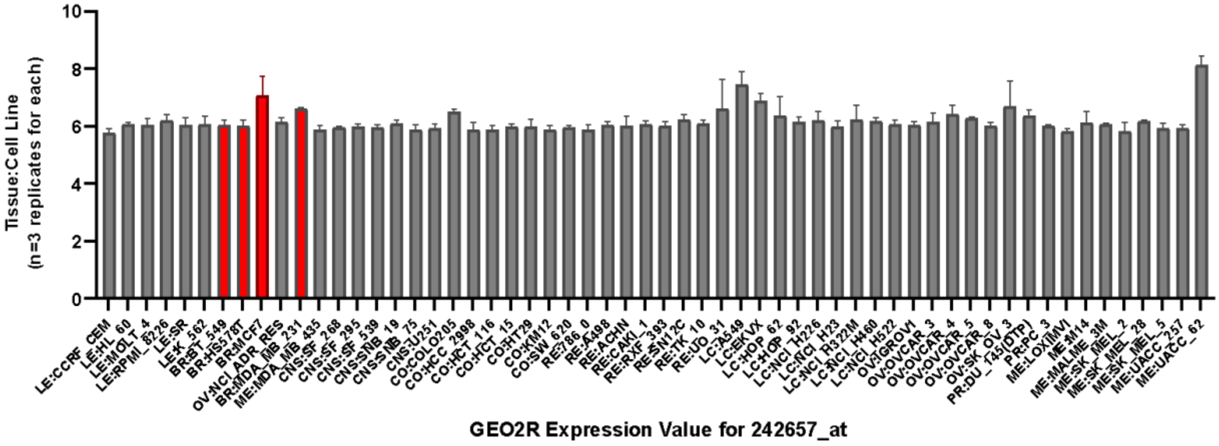


c

| Probe ID | Probe Number | Score | Start | End | Qsize | Identity | Chr | Str | Start | End | Span | Probe Sequence |
|---|---|---|---|---|---|---|---|---|---|---|---|---|
| 242657_at | 1 | 25 | 1 | 25 | 25 | 100.00% | chr17 | + | 38601999 | 38602023 | 25 | GCAAGGGCCGAGCCCTTGGGTTCGC |
| 242657_at | 2 | 25 | 1 | 25 | 25 | 100.00% | chr17 | + | 38602015 | 38602039 | 25 | ATCCTCCACCAGCCAGAGGGTATCT |
| 242657_at | 3 | 25 | 1 | 25 | 25 | 100.00% | chr17 | + | 38602030 | 38602054 | 25 | TAAGGATGGCTGGTAGCCTCCTCCT |
| 242657_at | 4 | 25 | 1 | 25 | 25 | 100.00% | chr17 | + | 38602059 | 38602083 | 25 | CTCCTCTCTCAGCACTTTGAATGCA |
| 242657_at | 5 | 25 | 1 | 25 | 25 | 100.00% | chr17 | + | 38602093 | 38602117 | 25 | GGCTGGGTAGGAATACCTCATCCTC |
| 242657_at | 5 | 20 | 4 | 25 | 25 | 95.50% | chr8 | - | 40966301 | 40966322 | 22 | GGCTGGGTAGGAATACCTCATCCTC |
| 242657_at | 6 | 25 | 1 | 25 | 25 | 100.00% | chr17 | + | 38602113 | 38602137 | 25 | ATACCTCATCCTCTGGTTGGGTAGG |
| 242657_at | 7 | 25 | 1 | 25 | 25 | 100.00% | chr17 | + | 38602158 | 38602182 | 25 | GGGTAGGAATACTCCATCCTGTAGT |
| 242657_at | 8 | 25 | 1 | 25 | 25 | 100.00% | chr17 | + | 38602170 | 38602194 | 25 | GTGGTTGTCCTCTGGATATAATGCA |
| 242657_at | 9 | 25 | 1 | 25 | 25 | 100.00% | chr17 | + | 38602188 | 38602212 | 25 | GATATAATGCAAACTGGCCTCCAGG |
| 242657_at | 10 | 25 | 1 | 25 | 25 | 100.00% | chr17 | + | 38602266 | 38602290 | 25 | TGGGTTCGCGGGGTACAACAGAGTC |
| 242657_at | 11 | 25 | 1 | 25 | 25 | 100.00% | chr17 | + | 38602280 | 38602304 | 25 | CAACAGAGTCACGTTCCCTTTGAAT |

**Figure S7. a.** The UCSC Genome Browser view displays the genomic location of the Affymetrix Human Genome U133 Plus 2.0 probes corresponding to both full-length and intronic regions of IGFBP4. The view also includes Affymetrix Human Exon Array extended probes, offering additional evidence for probe coverage within the region of interest, **b**. GEO2R expression data for *IGFBP4* EI isoform. 242657_at probe set maps to *IGFBP4* intronic region. Red bars indicate breast cancer cell lines. The data was retrieved from the NCI-60 Cancer Cell Line data (GSE32474 GPL570 platform), **c**. BLAT result showing the specificity of the 242657_at probe set for the intronic IGFBP4 region. Probe sequences were retrieved using the BiocManager package in R, and their specificity was confirmed using the UCSC BLAT.

a

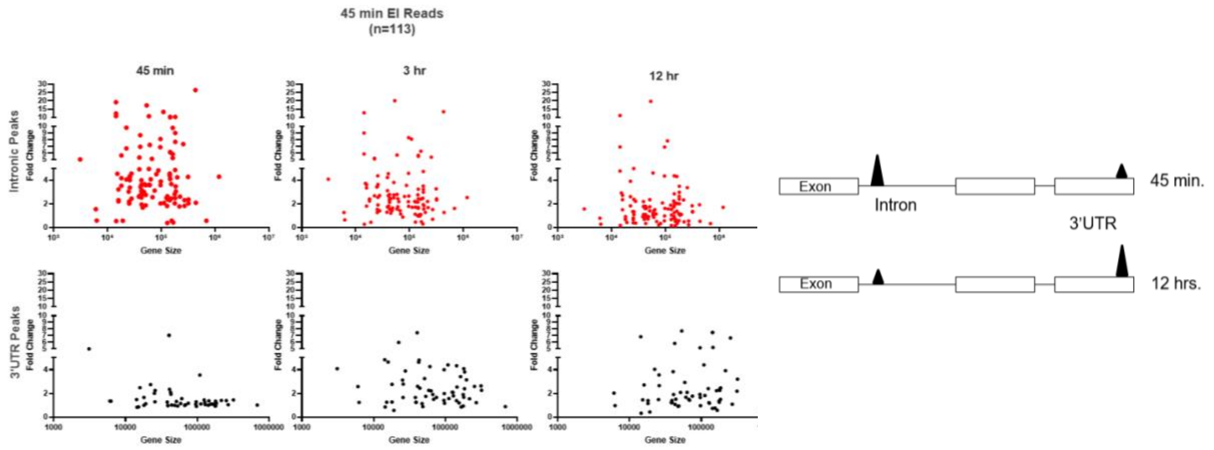


b

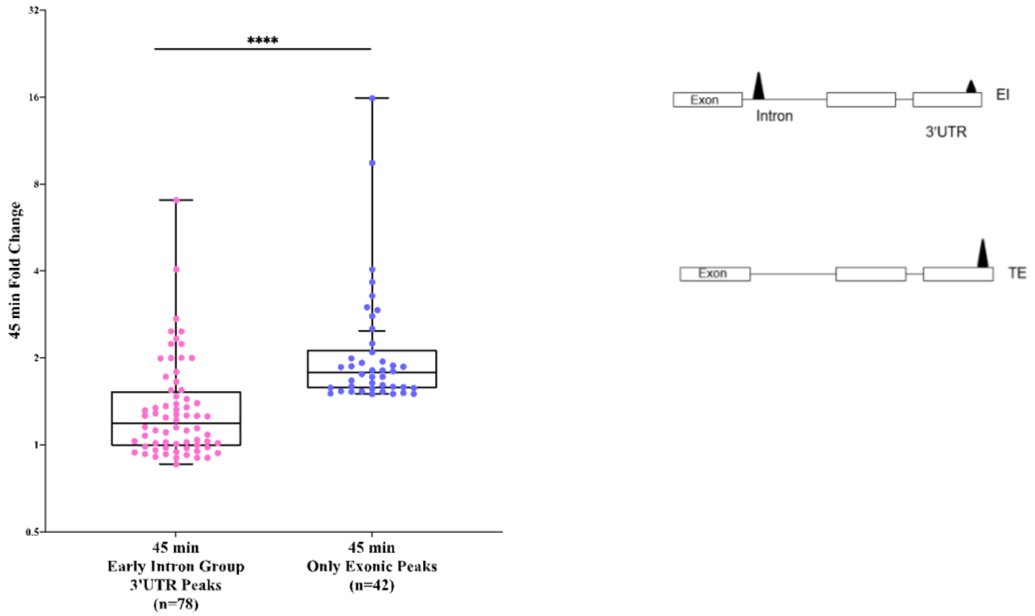


**Figure S8**. **a**. 45 min. EI genes (detected only by 3'-seq). Expression detected from early intronic regions (red) compared to corresponding 3'UTRs (black). The x-axis denotes gene sizes; the y-axis represents expression fold changes, **b**. Overall upregulation of full-length isoforms of 3'seq-only detected EI group genes (pink, top drawing) compared with genes that had only 3'UTR reads (blue, bottom drawing). Unpaired t-test (**** <0.0001).

a

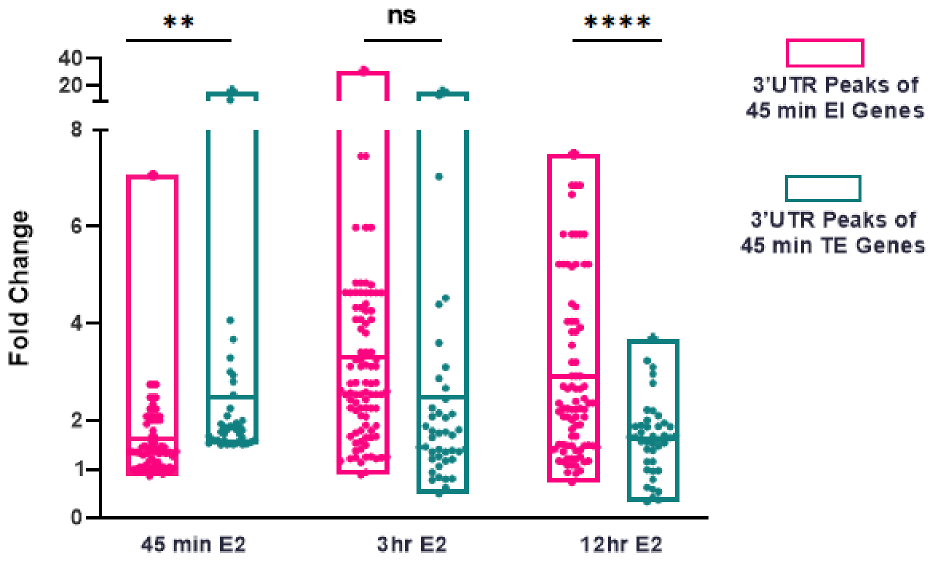


b

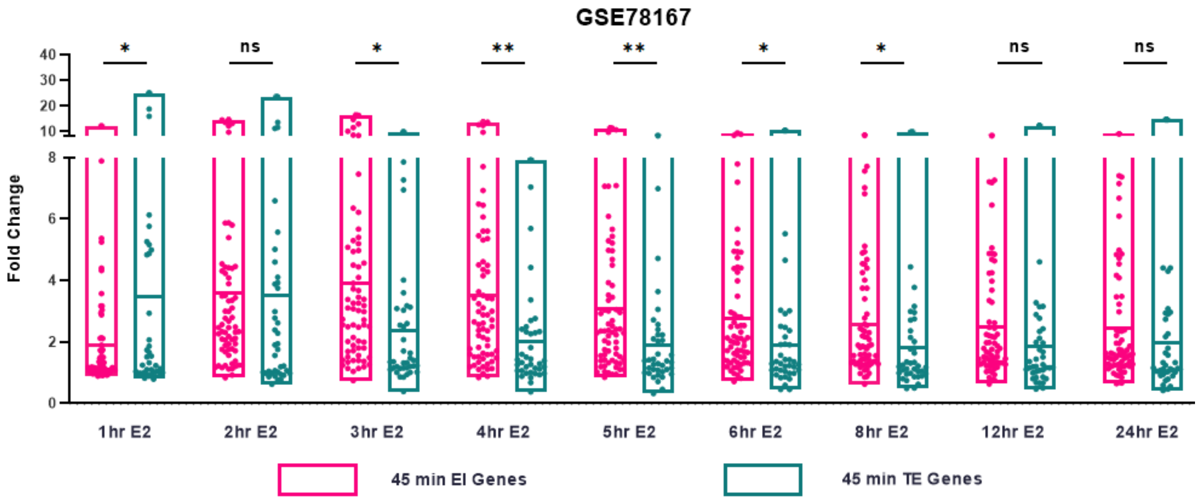


**Figure S9. a.** Expression based on terminal exon reads for the EI (pink) genes compared to TE (green) genes at 45 minutes of E2 treatment. Expression of this set of genes was followed for 3 hours and 12 hours, **b**. The overall expression of 45 min. EI genes compared to TE genes (RNA seq. data of GSE78167, MCF7 cells 10 nM E2 treatment). Statistical significance between EI and TE gene groups at each time point was determined using a two-sample student's t-test, ns (not significant, $p > 0.05$), * ($p < 0.05$), ** ($p < 0.01$), and **** ($p < 0.0001$).

a

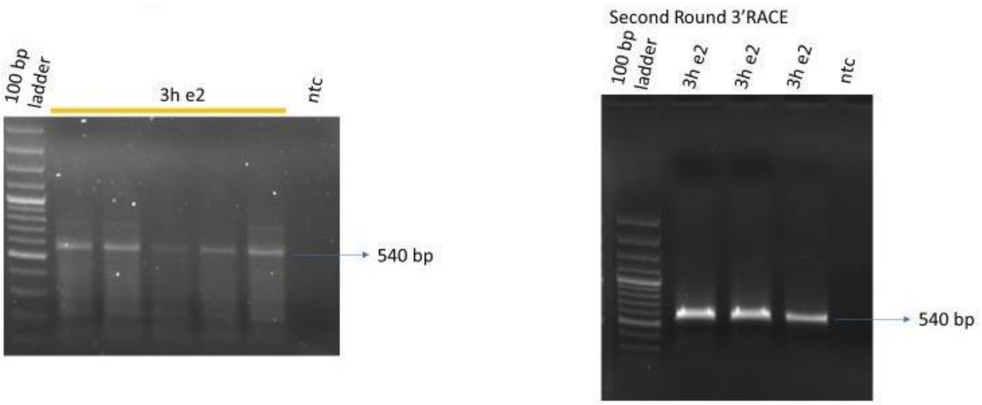


b

Forward Primer: 5' – GCAATGGGTCCGGCT – 3'

GCAATGGGTCCGGCTGTTTTCAACGGCAAAGATTCCCCGAAGTGCCAACAACTGGCAAAGAATAACCTGTTGGCCCTGCCGCGACCATCG
GCTTTAGGTAGCTCTGCCTGTCCTGGCCGTCCTGGGGATCCTGCTCTTCTTTGGGCCAAGGCCAGAGGGGGCATGGGAGCTGCTGGGCTG
GGGCCGCTGAGCCTCCTGGAGTCTGATGTGGCTTCCATGAGAGTCAGGCCAGGACCACAGGTGCCTCCTGAGTCCGTGGGTCTGGGCCC
AGGCCCTGGTGCTCTATGAGTGACACTTGGGAGTTACCGTATGTGCCAAGGAGCAGTGAGACACCTGTGGTAATGGCGAGTGACCTGTC
CTATGGGCTGCACAGCTGGAGGATGGTAGAATTTTGGAGCAATAGTGGGCAGTGTGTTGTTGGGGAAGAACATGGGCTTCTGAATCAGA
AAAATATGAATTCAAAACCTACCGCTGTCTCTTGTTAGTCAGGGGACCTTGGCCAAGTCTCAATAAATGTTTCTTCATCTGGA

Anchor Primer: 5'- TCCAGATGAAGAAACATTTATTGTTTTTTTTTTTTTTTT -3'

c

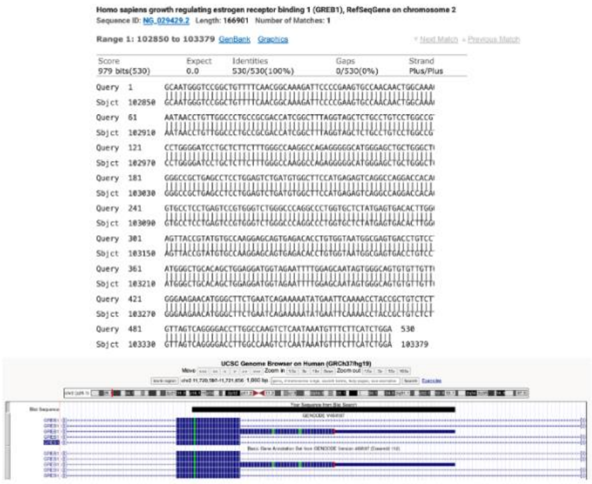

**Figure S10.** ***GREB1* 3'RACE a.** First and second round 3'RACE PCR products, **b.** Sequenced pGEM-T vector that harbored the cloned 3'RACE PCR product, **c**. cloned RACE product (black) in alignment with *GREB1* gene structure.

a

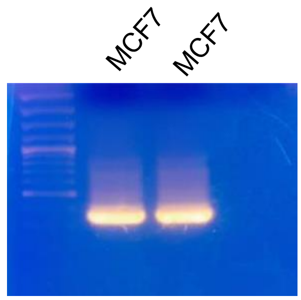


b

Forward Primer: 5’- AAAGTAATGTTGTTAAATGGGGCGA -3’

AAAGTAATGTTGTTAAATGGGGCGAAATAAGAAGTTAAATTTGAAAGTGAATGTGTTCAAATGAAAGGTTTGATAATTGCATC
TGTTACTACTTAGTTTCATAGGCTTTAATTCTAGTATGCATTAAATATTGGGCAAAATTGCACTTGACTAATTTTTTGAAGAAAA
GTAATTTATTCTGTCAAGAAATAAAAAAATAGGCTTTGTGTATGGTTAAACTGTAAATCTTATGTTTACAAAATACTGTAATTTT
CAGGAAATCACTGTATTAGGAATGTGCAATGACTTATATAAATAAAAGCCATTTTTAAAACTGAAAAAAAAAAAAAAAAAG

Anchor Primer: 5’- GACCACGCGATCGATTGACTTTTTTTTTTTTTTTTTV -3’

c

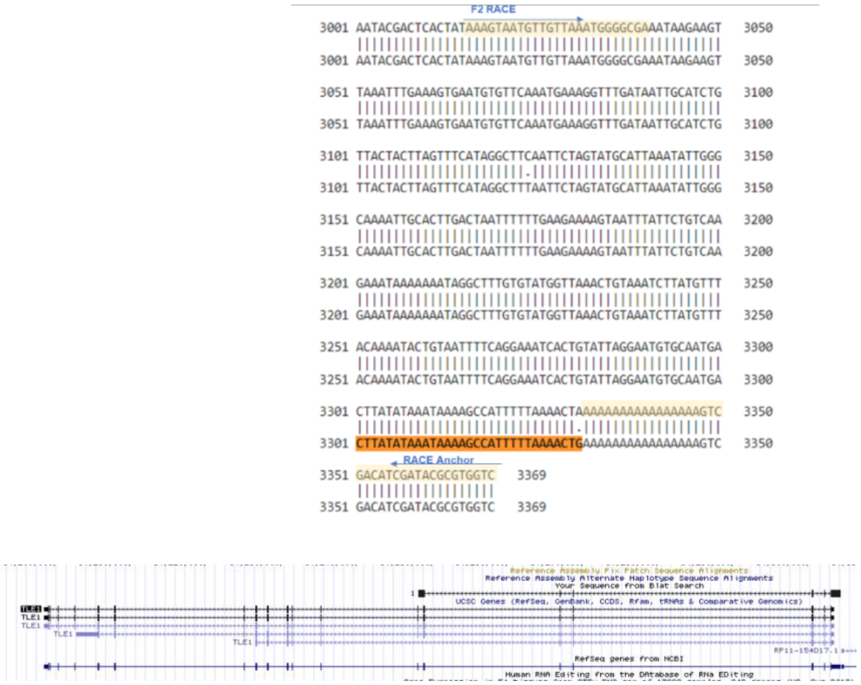


**Figure S11. *TLE1* 3’RACE, a.** 3’RACE PCR products. **b.** Sequenced pGEM-T vector that harbored the cloned 3’RACE PCR product, **c.** cloned RACE product (black) in alignment with *TLE1* gene structure.

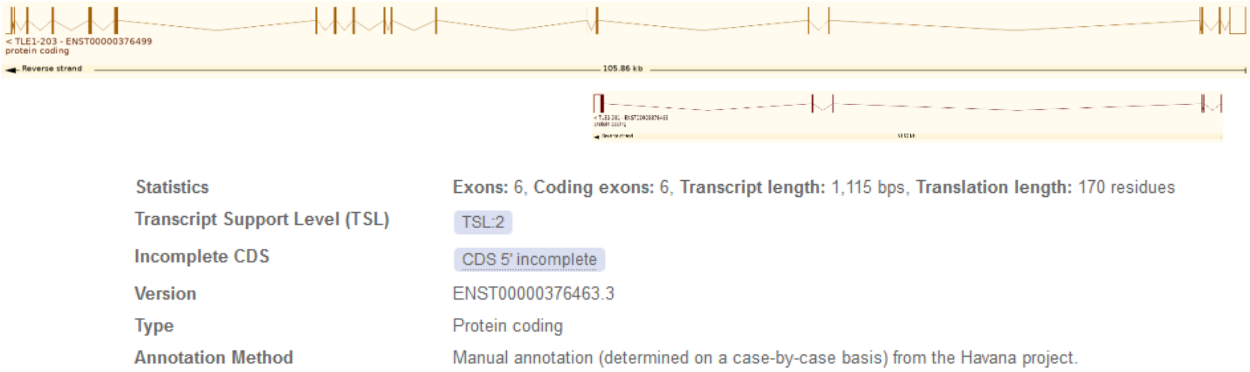


**Figure S12**. ENSEMBL entry for full-length TLE1 and ENST00000376463. Image taken from (http://www.ensembl.org/)

a

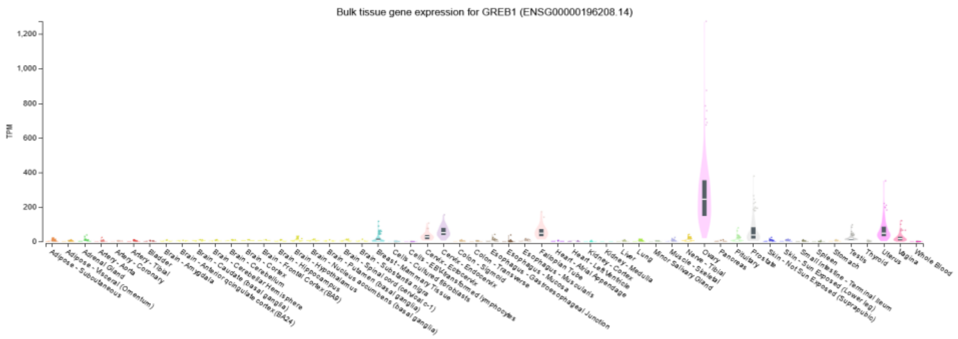


b

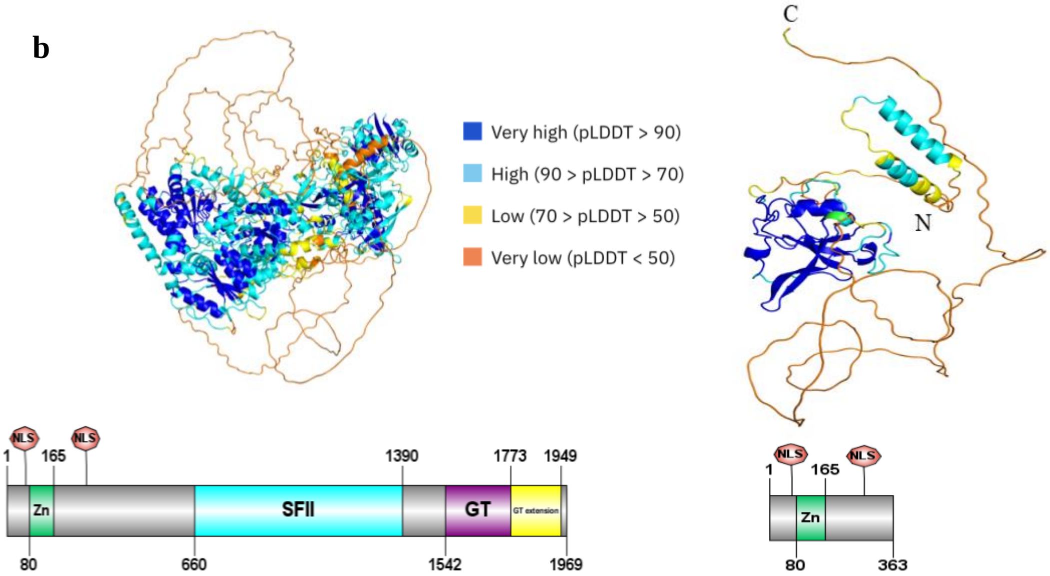


c

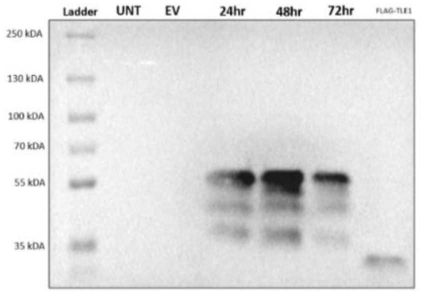


**Figure S13. GREB1 and C-term truncated GREB1 IPA**

**a.** GTEx expression of *GREB1* in normal tissues, **b.** AF2 protein models for the GREB1 proteins translated from the full-length and the IPA isoforms, colored according to AF2 plDDT confidence scores, protein domains found in GREB1 and GREB1 IPA are shown under the models. **c.** Western Blot analysis of truncated Flag-tagged GREB1 expression. Proteins were separated using a 5% stacking and 10% separating polyacrylamide gel. MCF7 cells were transiently transfected, and proteins were isolated at 24-, 48-, and 72-hours post-transfection. For each lane, 25 µg of protein was loaded. The expected molecular weights are 41 kDa for the GREB1-IPA. TLE1-IPA was loaded as a positive control for Flag-tag detection (25 kDa).

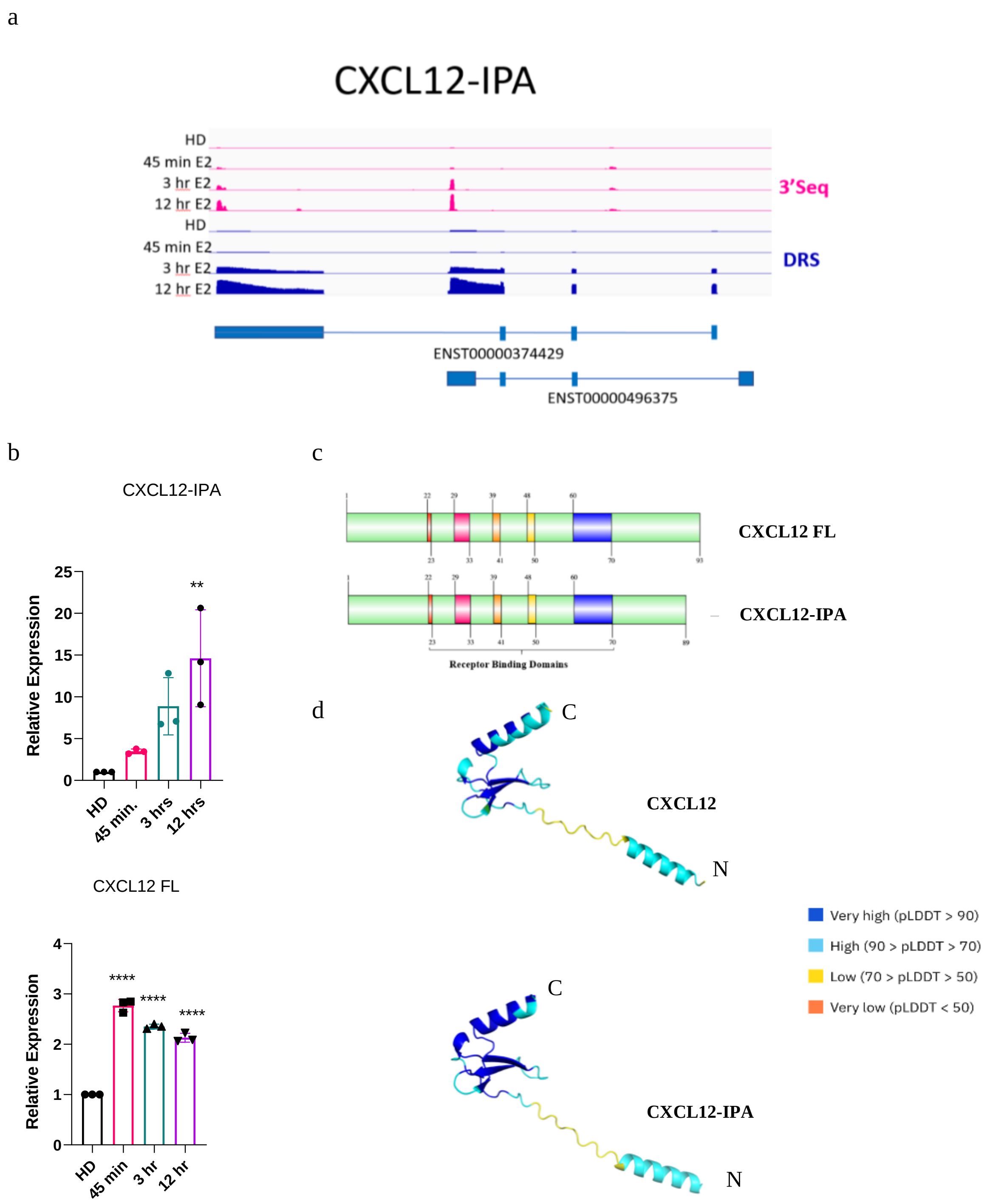


**Figure S14**. **a**. IGV display of *CXCL12* IPA and FL isoforms detected by DRS and 3'seq, **b.** RT-qPCR shows upregulation of IPA and full length (FL) isoforms for CXCL12 (n=3 independent E2 treatments, One-way ANOVA **p<0.005), **c**. CXCL12 (FL) and CXCL12-IPA protein domains, **d.** AF2 protein models for the CXCL12 proteins translated from the full-length and the IPA isoforms, colored according to AF2 plDDT confidence scores.
